# Bioinformatic analysis based genome-wide identification, characterization, diversification and regulatory transcription components of RNA silencing machinery genes in wheat (*Triticum aestivum* L.)

**DOI:** 10.1101/2020.05.21.108100

**Authors:** Zobaer Akond, Hafizur Rahman, Md. Asif Ahsan, Md. Parvez Mosharaf, Munirul Alam, Md. Nurul Haque Mollah

## Abstract

Dicer-Like (DCL), Argonaute (AGO), and RNA-dependent RNA polymerase (RDR) gene families are known as RNA silencing machinery genes or RNAi genes. They have important activities at post-transcriptional and chromatin modification levels. They regulate gene expression relating to different stresses, growth, and development in eukaryotes. A complete cycle of gene silencing is occurred by the collaboration of these three families. However, these gene families are not yet rigorously studied in the economically important wheat genome. Our bioinformatic analysis based genome-wide identification, characterization, diversification and regulatory components of these gene families identified 7 TaDCL, 39 TaAGO and 16 TaRDR genes from wheat genome against RNAi genes of *Arabidopsis thaliana*. Phylogenetic analysis of wheat genome with *Arabidopsis* and rice RNAi genes showed that TaDCL, TaAGO and TaRDR proteins are clustered into four, eight and four subgroups respectively. Domain, motif and exon-intron structure analyses showed that the TaDCL, TaAGO and TaRDR proteins conserve identical characteristics within groups while retain diverse differences between groups. GO annotations implied that a number of biological and molecular pathways are linked to RNAi mechanism in wheat. Gene networking between transcription factors and RNAi proteins indicates that ERF is the leading family linked to maximum RNAi genes followed by MIKC-MADS, C2H2, BBR-BPC, MYB, and Dof. *Cis*-regulatory elements associated to RNAi genes are predicted to act as regulatory components against various environmental conditions. Expressed sequence tag analysis showed that larger numbers of RNAi genes are expressed in different tissues and organs predicted to play roles for healthy plants and grains. Expression analysis of 7 TaDCL genes using qRT-PCR showed that only TaDCL3a and TaDCL3b had root specific significant expression (p-value<0.05) with no expression in leaf validated EST results. Besides, TaDCL3b and TaDCL4 significantly prompted in drought condition indicating their potential role in drought stress tolerance. Overall results would however help researchers for in-depth biological investigation of these RNAi genes in wheat crop improvement.

## 1. Introduction

Recently gene silencing technique has attracted many molecular biological scientists to develop transgene system in eukaryotic cell to solve complex biological problems and to make crop improvement by implementing the RNAi mechanisms. Plants and agricultural crops by nature establish some specific molecular mechanisms that help survive in diverse conditions in their life span. RNA silencing often known as RNA interference (RNAi) technique is one such mechanism, which is well-preserved in most multicellular eukaryotic groups and a unifying fact to define a sequence-specific RNA degradation process that controls sequence specific regulation of gene expression[1–3]. This technique was originally learned in plants and occurs broadly in eukaryotic organisms[4,5]. Eukaryotic organisms generate copious types of small RNAs[6]. It has been explored in previous studies that gene silencing or RNAi technique is triggered and maintained by double-stranded RNA(dsRNA) that provides increase to small RNA molecules[1,2,7–11]. There are two types of small RNA molecules nearly of size 20-30 nucleotides long ribo-nucleotides termed as microRNA (miRNA) and short interfering RNA (siRNA)[2,8,10,11]. microRNAs (miRNAs) are derived from self-complementary hairpin structures, whereas small-interfering RNAs (siRNAs) are resulting from double-stranded RNA (dsRNA) or hairpin precursors[12,13]. These minute size RNA particles play roles in controlling various biological and genetic processes in terms of growth and development, creation of heterochromatic, retaining genome stability by means of regulation of gene expression and suppression of transposon movement, anti-pathogen and fungal defense[1,3,10,11,14–16]. These small RNAs are mainly linked to both post-transcriptional gene silencing and triggering chromatic modifications[10,17]. In plants and crops, the production and function of these miRNAs and siRNAs are largely rely on three main protein families known as Dicer-like(DCL), Argonaute(AGO) and RNA-dependent RNA polymerase(RDR)[8,10]. A complete cycle of RNA silencing takes three steps: initiation, maintenance and signal amplification. The beginning of gene silencing requires the production of double-stranded RNA (dsRNA). dsRNAs that are created from plant-encoded RNA-dependent RNA polymerases(RDRs) on a variety of RNA templates[3]. The other processes of creation of dsRNA include bidirectional transcription of DNA, self-complementary RNA fold backs [7]. The complementary dsRNA are then processed by the act of RNaseIII-type enzymes called Dicer (DCR) in animals or Dicer-like (DCL) in plants which then alter these dsRNA into tiny RNAs of 19-31 nucleotides(nt) in size siRNA/miRNA[3,7,10]. Afterward, one strand of these siRNAs is attached to AGO protein containing complexes termed as RISC (RNA-induced silencing complex). RISC possesses endonuclease ability to operate cleavage activity of target mRNAs or DNAs that are complementary/homologous to siRNA/miRNA by means of AGO’s RNaseH-type enzyme property for RNA degradation, translational inhibition, or heterochromatin formation [1,3,7,8,10,18]. siRNAs are also predicted for transcriptional gene silencing by implementing RNA-directed DNA methylation or chromatic remodeling to regulatory sequences of target genes[7,10]. At the signal augmentation step, RDR enzymes are likely for synthesis of dsRNAs from ssRNA templates to start a new cycle of RNA silencing [8,19]

DCL, AGO and RDR protein families are however considered as the chief components of gene silencing or RNAi cycle. DCL proteins are known as the important protein families among the other RNA silencing machinery genes for small RNA biogenesis routes that transform the long dsRNAs into mature miRNAs and siRNAs[10,11]. DCL families are identified and characterized to possess six conserved domains, viz., dead box, helicase, dicer dimerization, PAZ (Piwi Argonaut and Zwille), RNase III, and dsRNA binding domain (dsRBD)[7,8,10,11]. Although one or two domains are assumed to absent in the lower type eukaryotic organisms[11,20]. The helicase domain is termed to identify the single strand terminal loop structure of primary miRNA(pre-miRNA) and limits the proportion of substrate cleavage [11,21]. The dicer dimerization domain is engaged in target selection of small RNA handling [22]; the PAZ and RNaseIII domains contribute in dsRNA substrate cleavage[11,20]. Surprisingly, the distance between PAZ and the cleavage site in the RNaseIII domain decides the size of matured small RNAs[11,23]. DCL however contains a minor set of gene family in upper plants, insects, protozoa, and some fungi, while only one Dicer protein is present in vertebrates, nematodes, *Schizosaccharomyces pombe*, and green alga *Chlamydomonas reinhardtii*[8,20].

AGOs are also known as the special kind of protein families in RNA silencing mechanism located at multi-protein type RISC that direct to cleave the target mRNAs, which are complementary sequence of siRNAs. AGO proteins particularly in larger nearly 90-100kDa characterized by the presence of four functional domains or amino acid motifs, viz., ArgoN/Argo-L, C-terminal PIWI(P-element induced wimpy testis),PAZ and MID[6,24,25]. A chief PAZ domain possesses a particular binding pocket that binds siRNAs over the 3’ end of the target mRNA[10,24] whereas a PIWI binds siRNAs through the 5’ end of the target mRNA which holds the complete homology characteristics like RNaseH to cleave target mRNAs[1,10,26]. MID domains are placed between the PAZ and the PIWI domains. A very simple pocket characteristic of such MID domain particularly binds the 5’ phosphate of the siRNAs and then anchors the siRNAs onto AGO proteins [10,26]. In eukaryotic species, minimum three subclasses of AGO proteins have been recognized[7]. These consists of AGO subfamily which is exist in plants, animals and yeasts; the Piwi and worm-specific AGO(WAGO) subgroups are observed in animals and *C. elegans* respectively[7,27–29]. However, all members of the subgroup of AGO and Piwi conserve the particular DDH catalytic residues but absent in WAGO. The expression of Piwi subgroup proteins is limited to human germ cell and in rat and some mammals. Although this cluster of AGO proteins have not been identified in any plant species but in recent times a novel type of Argonaute, OsMEL1 has been identified in rice which is linked to male meiosis [7,30].

The third type of RNA silencing machinery gene is the RDR protein. This kind of protein family is exist and highly essential for RNAi mechanism in fungi, nematodes and plants, but have not been identified in insects or vertebrates[31]. RDR proteins were separated first from tomato[32]. RDR proteins are critically important for start and increase of silencing process and these protein families contain a well-preserved sequence motif like the catalytic β’ subunit of DNA dependent RNA polymerases[7,33]. This gene family, however, possesses a RNA-dependent RNA polymerase (RdRP) domain, help form the dsRNA from single-stranded RNAs (ssRNAs) to start a new cycle of RNA silencing [7,34].

Recent study suggests that there are numerous copies of DCL, AGO and RDR genes are present in plants and animals[7,10]. Different study on a dozen of crops plant has been made to learn about the role of these RNAi gene families. All members of each of these gene families take part in diverse roles in RNA silencing pathway. A model plant in plant biology, for instance, *Arabidopsis thaliana* possesses 4 DCL (AtDCL1-AtDCL4) that definitely generate diverse kinds and sizes of small RNAs [18]. To activate various important biological functions in eukaryotic cell, different small RNAs produced in cell with diverse functions but they all are the associate members of DCL, AGO and RDR gene families. However, in *Arabidopsis thaliana* genome, 4 AtDCLs, 10 AtAGOs and 6 AtRDRs genes were identified [6,7,10]. 4 NbDCL, 9NbAGO and 3 NbRDR RNAi genes were identified in tobaco(*Nicotiana benthamin*)[35]. Rice (*Oryza sativa*), a crop of monocot group, possesses 8 OsDCLs, 19 OsAGOs and 5 OsRDRs genes, in which OsAGO2 gene exhibited particular up-regulation in response to salt and drought [7,32]. Also, in tomato (*Solanum lycopersicum*), 7 SlDCLs, 15 SlAGOs and 6 SlRDRs genes were identified [36]. Five ZmDCLs, 18 ZmAGOs and 5 ZmRDRs genes were recognized in maize(*Zea mays*) genome [10]. There are 4 VvDCLs, 13 VvAGOs and 5 VvRDRs genes were detected in grapevine (*Vitis vinifera*)[37]. Likewise, 5, 7, and 8 CsDCLs, CsAGOs, and CsRDRs, respectively have been identified in cucumber (*Cucumis sativus*) [38]. On the other hand, the genome of allopolyploid species of *Brassica napus* contain 8 BnDCLs, 27 BnAGOs, and 16 BnRDRs [8,39]. Overall, 4 CaDCLs, 12 CaAGOs and 6 CaRDRs genes have been identified in pepper (*Capsicum Annuum*) [1].In coffee (*Coffea canephora*) 9CcDCLs, 11CcAGOs and 8 CcRDRs were identified for genome-wide study by means of RNA guided pathways in 2017[2]. Foxtail millet (*Setaria italica*) possesses 8 SiDCL, 19 SiAGO and 11 SiRDR genes. A very recent study suggests that sweet orange (*Citrus Sinensis*) genome possesses 5CsDCLs 13 CsAGOs, and 7 CsRDR RNAi genes [40]. However, successive investigation on RNAi genes in various important crop and fruit plants showed considerable divergence and have indispensable role in different genomic functions. In *A. thaliana,* AtDCL1 principally play role for miRNA biogenesis while AtDCL2, AtDCL3, and AtDCL4 mediate siRNA processing[10,11]. Besides, AtDCL3 and AtAGO4 are essential for RNA-directed DNA methylation of the FWA transgene, which is associated to histone H3 lysine 9 (H3K9) methylation[10,11]. Though AtDCL2 produces siRNAs responsible to create defensive mechanism against viral infection and by nature the creation of siRNAs are tend to happen from *cis*-acting antisense transcript but AtDCL4 regulate the vegetative stage transformation by the production of siRNAs from *trans*-acting transcript[10,41,42]. Surprisingly,AtDCL3 selects short dsRNAs but AtDCL4 cleaves long dsRNA substrate s[11]. Many other functions of DCL genes in plants such as AtDCL1 and AtDCL3 proteins contribute in promoting flowering[43].

On the other hand, AGO gene families act as the chief RNA-mediated gene silencing techniques and reported that they play noteworthy performance in growth and development in plants[24,44,45]. AtAGO1 is associated with the transgene silencing pathways[46] and AtAGO4 with the epigenetic silencing [47]. AtAGO7 and AtAGO10 however effect the growth [48] and meristem maintenance [49]. Other AtAGOs however have some important characteristics in gene silencing pathways. Earlier investigations also suggest that the RDR genes are biologically prompt in RNAi mechanism such as co-suppression, protect pathogen invasion, chromatin modification, and post-transcriptional gene silencing activities in plants, viz., *Arabidopsis*, maize [50–54].

Wheat (*Triticum aestivum*), a global common staple food as well as the second most-produced cereal crops after maize in the world (http://www.fao.org/worldfoodsituation/csdb/en/). It is also the high vital source of carbohydrates and is the leading source of vegetal protein in human food. There is little systemic thorough study has been made so far regarding RNA silencing machinery genes for this crop. The current study was carried out to obtain comprehensive bioinformatic understanding in terms of *in-silico* identification, characterization, diversity survey, regulatory components, and expression sequence tag(EST) analysis of all members of DCL, AGO and RDR gene families in wheat. Finally, wet-lab experimental expression analysis was investigated for the identified 7 wheat DCL proteins in the two wheat plant organs (root and leaf) and in drought condition as well. Outcomes obtained from this work will offer fundamental source of genomic information for these RNAi genes and will certainly help plant molecular researchers for insights into the possible roles of these genes in plant growth, development, and performances against various biotic and abiotic stressors. The systemic schemes of this whole study in this article however have been demonstrated briefly in Fig. 1.

**Fig 1.**
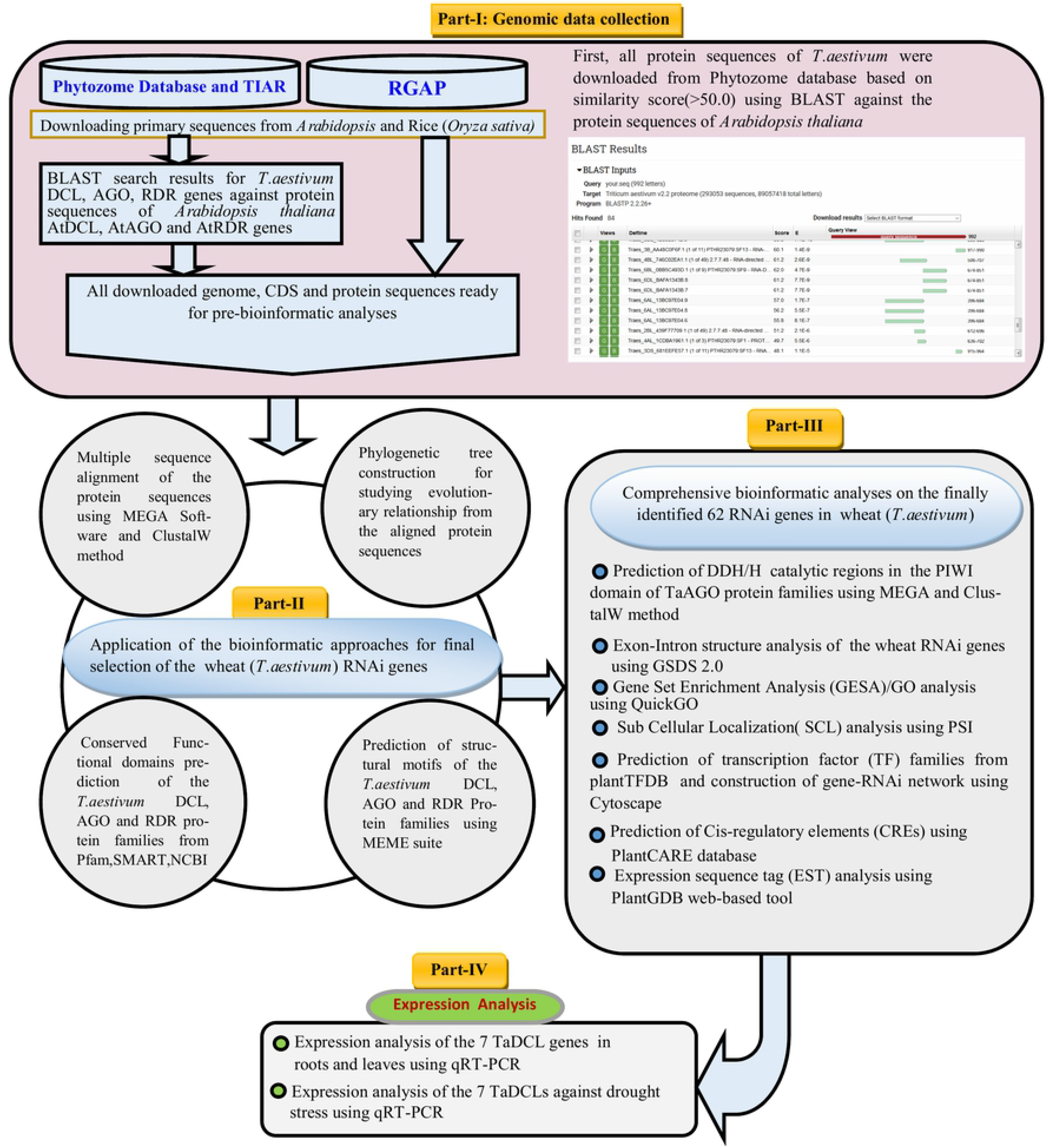
The flow diagram of the complete work in this study.

## 2. Materials and methods

### 2.1. Data sources of RNAi genes

Protein sequences of *Arabidopsis* and rice (*Oryza sativa*) DCL, AGO and RDR were obtained from The Arabidopsis Information Resource (TAIR) (http://www.arabidopsis.org) and Rice Genome Annotation Project (RGAP) (http://rice.plantbiology.msu.edu/) respectively following the profile Hidden Markov Model (profile HMM). Then these sequences were used as query sequence to search using Basic Local Alignment Search Tool (BLAST-P) program (p-value=0.001) against *T. aestivum* genome from the plant genome Phytozome database (http://phytozome.jgi.doe.gov/pz/portal.html). Paralogous protein sequences from the wheat (*Triticum aestivum*) genome were collected by considering the threshold value (≥50) and significant E-values. Only the primary transcripts were selected to avoid the redundancy of the sequences in the analysis. All retrieved sequences were searched for conserved functional domains using the Pfam (http://pfam.sanger.ac.uk/), web-based Simple Modular Architecture Research Tool (SMART)(http://smart.embl-heidelberg.de/) and the National Center for Biotechnology Information Conserved Domain Database (NCBI-CDD: http://www.ncbi.nlm.nih.gov/Structure/cdd/wrpsb.cgi). Some basic information of these genes such as accession number, genomic location, gene length, and encoded protein length were downloaded from Phytozome wheat genome database. All newly identified genes from wheat genome were designated as TaDCL, TaAGO and TaRDR following taxonomy based on phylogenetic relationship of the same family members of the *Arabidopsis thaliana* genes as termed earlier investigations. Prediction of the molecular weight of each gene member of each DCL, AGO and RDR group was predicted from web-based tool ExPASyComputepI/Mwtool (http://au.expasy.org/tools/pitool.html).

### 2.2 Bioinformatic analysis approaches

Several bioinformatic analysis approaches including sequence alignment and phylogenetic tree construction, prediction of functional domain and motif structure of the proteins, exon-intron structure of the RNAi genes, gene ontology (GO) analysis, prediction of subcellular location, regulatory network among the gene transcription factors and wheat RNAi genes, *cis*-regulatory element(CRE) analysis, express sequence tag (EST) analysis were carried out for comprehensive genome-wide identification, characterization, diversification and regulatory transcription components of wheat RNA silencing machinery genes. These approaches are described in the following sub-sections 2.2.1-2.2.8.

#### 2.2.1 Sequence Alignment and Phylogenetic Tree Analysis

Protein sequences of DCL, AGO and RDR genes belong to respective *Arabidopsis thaliana*, rice and wheat were used for multiple sequence alignment using ClustalW method [55]. After that, to study the evolutionary relationship, three phylogenetic trees were constructed of the corresponding 7 TaDCLs, 39TaAGOs and 16 TaRDRs using MEGA7: Molecular Evolutionary Genetics Analysis software[56] with Neighbor-Joining (NJ) method. Phylogeny test was performed by following bootstrap method with 1000 replications to measure statistical support for nodes[57] and the evolutionary distances were computed using the p-distance substitution method with complete deletion of the gaps/missing data[56].

#### 2.2.2 Conserved functional domain and motif structure analysis

Conserved functional domains for each of the TaDCL, TaAGO and TaRDR protein family were predicted from the three well-established genomic online databases, viz., Pfam, SMART and NCB-CDD by using their corresponding protein sequences to search similar conserved functional domains contained by RNAi genes of *Arabidopsis Thaliana* AtDCL, AtAGO and AtRDR. Higher numbers of functional domain possessing TaDCL, TaAGO and TaRDR genes corresponding to that of AtDCLs, AtAGOs and AtRDRs genes were selected. Six functional domains,viz.,DEAD/ResIII, Helicase-C, Dicer-Dimer, PAZ, RNase III and DSRM were taken into consideration to select the potential DCL group genes(TaDCLs) in *Triticum aestivum*. The AGO gene families in wheat that contain Argo-N/Argo-L, PAZ, MID and PIWI important functional domains are termed as AGO class genes in wheat. TaRDR gene families were also listed for the presence of RdRP and RRM functional domains for protein structure.

In order to identify the most potential conserved metal-chelating catalytic triad regions or residues in the PIWI domain, i.e., aspartate, aspartate and histidine (DDH)[7] as well as histidine at 798 positions (H798), TaAGO protein families were considered. In addition, the two conserved motifs, viz, DLDGD and DFDGD in the RdRp domain of TaRDR protein groups were also detected. The alignment profiles for this identification were performed using ClustalW program in MEGA and GENEDOC software tool. All paralog AtAGO and AtRDR protein families were considered during alignment (Fig. 2). Moreover, the conserved structural motif differences among DCL, AGO and RDR gene groups in *T. aestivum* were projected using a web-based tool called Multiple Expectation Maximization for Motif Elicitation (MEME-Suite)[58]. MEME analysis was carried out by specifying the parameters: (i) optimum motif width as ≥6 and ≤50; (ii) maximum 20 motifs.

**Fig.2.**
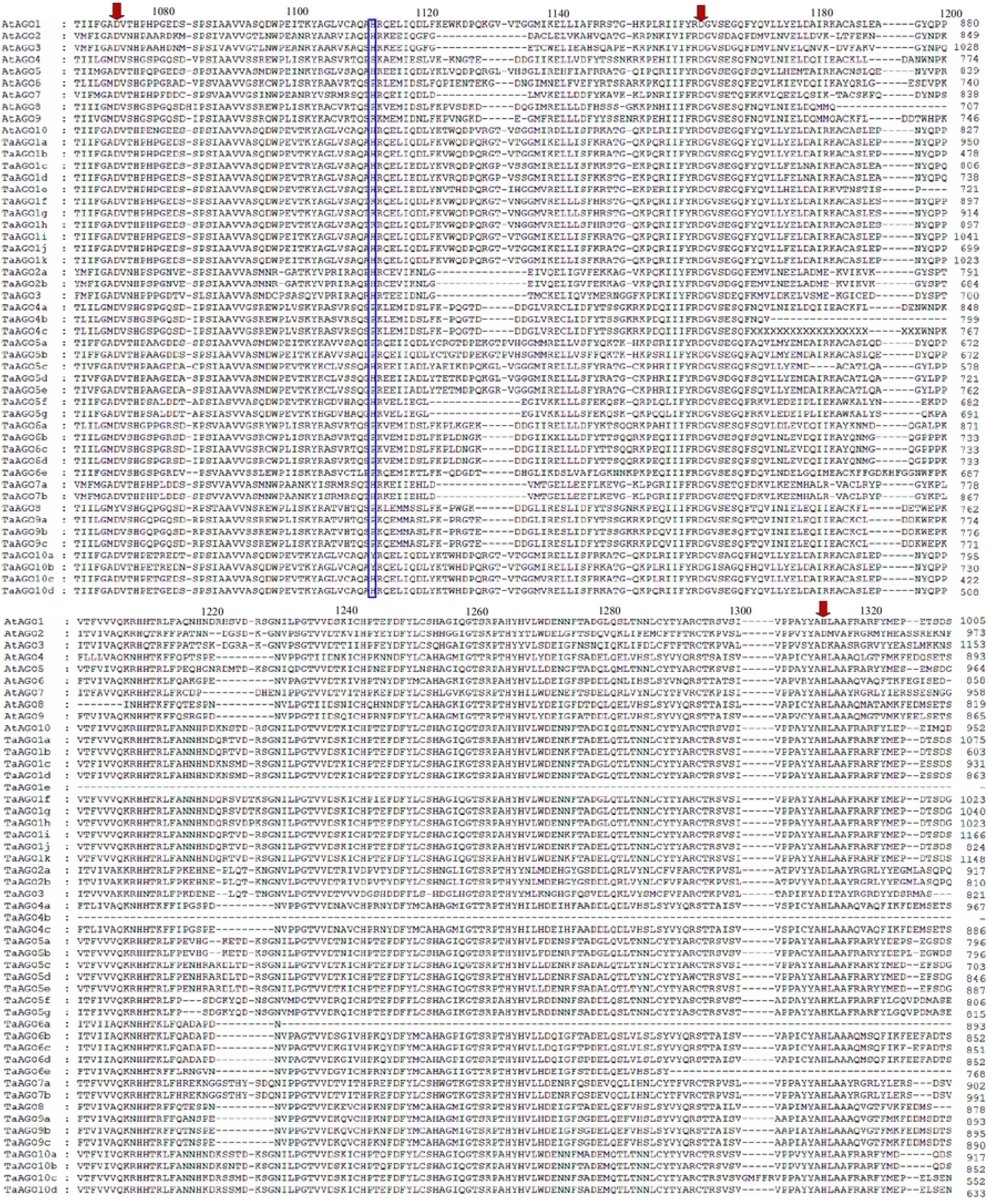
Alignment profile of PIWI domain amino acids of wheat (*T.aestivum*) AGO proteins. The protein sequences were aligned using MEGA 7.0. The conserved Asp, Asp and His (DDH) traid residues corresponding to D760, D845, and H986 of *Arabidopsis* AGO1 are indicated with downward arrows, whereas the conserved H residue corresponding to H798 of *Arabidopsis* AGO1 is boxed with blue color. Amino acid positions corresponding to each protein are indicated at the end of each line.

#### 2.2.3. Exon-intron structure analysis of the TaDCL, TaAGO and TaRDR gene families

The organization of the exon-intron structure of 7 TaDCLs, 39 TaAGOs and 16 TaRDRs genes were measured using the online-based Gene Structure Display Server (GSDS 2.0 : http://gsds.cbi.pku.edu.cn/index.php) by taking their respective complete coding sequences (CDS) and DNA/genomic sequences downloaded from Phytozome database. *Arabidopsis thaliana* 4 AtDCLs, 10AtAGOs, and 6AtRDRs genes were also included to observe the similarity study.

#### 2.2.4. Gene ontology and sub-cellular localization analysis

Gene ontology (GO) enrichment analysis was performed using the web-based PlantTFDB v4.0[59] and QuickGO (https://www.ebi.ac.uk/QuickGO) tool in order to investigate the particular involvement of the RNAi genes of *T. aestivum* in terms of biological function (BP), molecular function(MF) and in certain cellular component(CC). Fisher’s exact test was considered to determine respective p-values and Benjamin-Hochberg’s adjustment correction. Enrichment outcomes with adjusted p < 0.05 were considered as statistically important. Furthermore, a web-based tool called PSI (Plant Subcellular localization Investigative Predictor) [60] for predicting the subcellular location of the 62 RNA silencing genes of *Triticum aestivum* and open source software R-3.5.2 [61] were used. A protein was considered locating in the certain cellular location if p < 0.00001 in the resulting output of the tool.

#### 2.2.5. Identification and characterization of regulatory TFs and RNAi genes in *T.aestivum*

To identify and study the associated regulatory transcription factor (TF) families and their degree of relationship with the identified RNAi genes of *T. aestivum* were identified with the help of plantTFDB (http://planttfdb.cbi.pku.edu.cn//).

#### 2.2.6. Construction of gene regulatory network between TFs and *T.aestivum* RNAi genes

To explore the interacting regulatory association between TFs and RNAi genes, all identified TF families and predicted *T.aestivum* RNAi genes were used to construct a whole regulatory gene network and sub-networks to find out the leading TF components and hub proteins by means of interaction network using Cytoscape 3.7.1[62].

#### 2.2.7. Promoter *cis*-acting element prediction of the RNAi genes in *T.aestivum*

*Cis*-acting elements are highly conserved and commonly related to regulate the transcription of a particular gene or gene families in conjunction with the respective TFs. For in-depth analysis of the *T.aestivum* RNAi genes, the *cis*-acting regulatory components in the promoter region were collected and analyzed using web-based prediction bioinformatic tool PlantCARE(http://bioinformatics.psb.ugent.be/webtools/plantcare/html/). The extracted bundle of *cis*-acting components were classified into five different groups based on their involvement in various actions such as stress responsive, light responsive, hormone responsive and other which have unknown functions.

#### 2.2.8. Expressed sequence tag (EST) analysis

*In-silico* EST investigation was performed for the identified wheat RNAi genes to predict their expression status in various important wheat plant tissues and organs using the web-based database PlantGDB (http://www.plantgdb.org/cgi-bin/blast/PlantGDB/). The DNA sequences of each TaDCL, TaAGO and TaRDR gene group were used to extract the EST information. The default parameter with e-value=1e−04 was set for blastn search. Finally, a heatmap was created to characterize the particular RNAi gene expression into various tissues and organs in *T.aestivum*.

### 2.3. RNA isolation and expression analysis of 7 TaDCL genes in *T. aestivum*

Complete RNA was taken out using Trizol reagent (TAKARA, Japan) by following the manufacturer’s guidelines. RNA was treated with DNaseI (TAKARA, Japan) and reverse-transcribed into cDNA using the PrimeScript RT reagent kit (TAKARA, Japan). The collected cDNAs were used for gene expression studies with quantitative real time PCR (qRT-PCR). The qRT-PCR was accomplished in StepOne Real-Time PCR System (Applied Biosystems, USA) using SYBER Premix Ex Taq reagents (TAKARA, Japan) following the program: 95◦C for 30s, 95◦C for 5s and 60◦C for 45s for 40 cycles. To normalize the sample variance, *T.aestivum* 18s gene was used as internal control. Relative gene expression values were calculated using the 2^−ΔΔ*Ct*^ method[63]. The primers used for gene expression analyses are listed in S1 Table. For the statistical analysis of the gene expression data, ANOVA was performed with SPSS software (Version19.0, IBM, USA). An important difference between mean values was assessed following DMRT (p < 0.05) approach.

## 3. Results and discussion

### 3.1. Identification and characterization of TaDCL, TaAGO and TaRDR genes

All downloaded protein sequences were undergone several analysis procedures for selecting the best possible RNAi genes in *T.aestivum* (Fig. 1). Then in total 62 RNAi gene set (7 TaDCL, 39 TaAGO and 16 TaRDR) were finally selected in wheat (*T. aestivum*) for subsequent bioinformatic analyses. By considering six functional domains, viz. DEAD/ResIII, Helicase-C, Dicer-dimer, PAZ, RNase III (Ribonuclease-3), and DSRM, 7 DCL loci were isolated and recognized in *T.aestivum* as TaDCLs using profile HMM with expect value threshold=0.01[64]. Conserved functional domains search of these 7 TaDCLs from the two databases(Pfam and NCBI-CDD) and SMART analysis reveals that almost all TaDCL protein families contain the said six domains (DEAD/ResIII, Helicase-C, Dicer-dimer, PAZ, RNase III and DSRM) which are termed as the full characteristics of plant DCL protein structure from the DCL family (class 3 RNase III family)[65].Though all TaDCLs contain DEAD/ResIII but this helicase type DEXH-box domain of endoribonuclease dicer was absent in TaDCL3b and TaDCL4. This domain typically contains the ATP-binding region[66]. Domain identification using SMART showed that the Dicer-dimer domain is absent but Pfam and NCBI-CDD database analysis showed the presence of this domain. This domain is found in members of the Dicer protein family, which functions in RNAi pathway and has weak dsRNA-binding action[64]. It facilitates heterodimerization of Dicer proteins with their corresponding protein partners[67]. Pfam analysis showed another domain of unknown function (DUF2838) in TaDCL1a. It is observed that all TaDCL protein families possess second DSRM domain shown by Pfam, SMART and NCBI-CDD databases that is entirely missed in non-plant DCLs, as described earlier [41]. The length of the TaDCL protein families however vary from 14743bp for TaDCL4 to 8347bp for TaDCL3d with their corresponding coding potential of 1392 and 1586 amino acids respectively (Table 1).

**Table 1.**
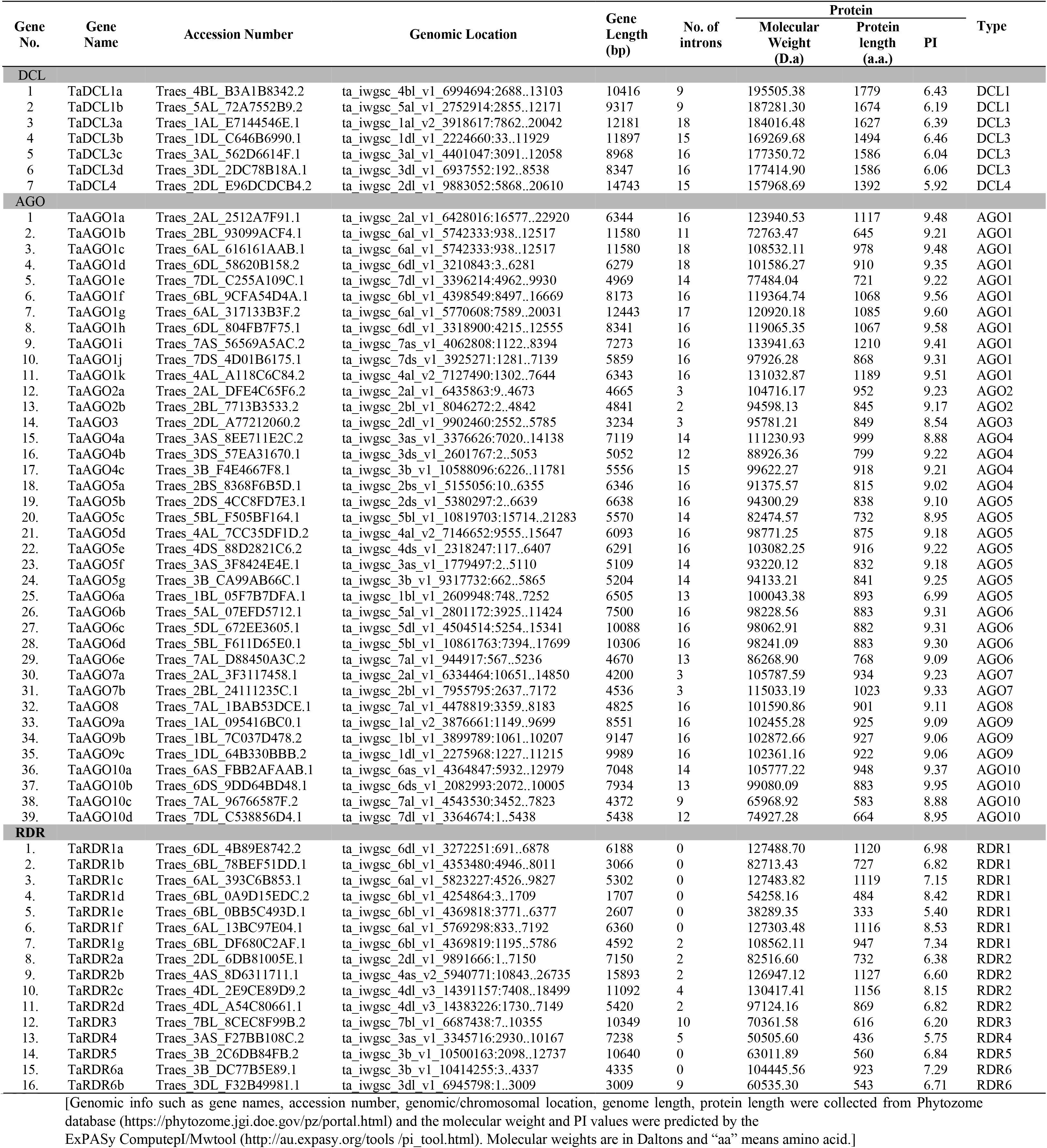
Basic genomic information about TaDCL, TaAGO and TaRDR protein families in wheat (*T.aestivum*).

Through HMM analysis for both conserved functional domains, viz., PAZ and PIWI using the downloaded protein sequences, 39 AGO RNAi protein families were identified finally for wheat (*T. aestivum*) genome. Conserved functional domain analysis using Pfam, SMART and NCBI-CDD showed that all identified putative TaAGO protein families are tend to contain an N-terminus PAZ domain and a C-terminus PIWI super family domain that are known as the main characteristics of plant AGO RNAi genes. It is also shown that other than these two important domains, wheat TaAGO proteins retain some other functional domains like *A. thaliana*, viz., ArgoN, ArgoL1, DUF1785, ArgoL2, ArgoMid. The length of the TaAGO protein families however varied from 12443bp for TaAGO1g to 3234bp for TaAGO3 with their corresponding coding potential of 1085 and 849 amino acids respectively (Table 1).

Besides, earlier investigations suggest that the PIWI domain presents complete homology to RNase H binds the siRNA 5’ end to the target RNA[26] and take part to cut target mRNAs that display sequence complementary to siRNA or miRNAs[68,69]. It is worth mentioning here that the three catalytic regions or residues in the PIWI domain such as aspartate(D), aspartate(D) and histidine(H) called DDH [7] are linked to the previous action and this triad was originally explored in *A.thaliana* AGO1, and a well-preserved histidine at position 798(H798) was also identified to be crucial for AGO1 for *in vitro* endonuclease function[69]. PIWI domains of all TaAGOs were aligned along with the paralogous 10 AtAGOs through multiple sequence alignment technique from their corresponding amino acid sequences using ClustalW program implemented in MEGA 7.0 (Fig. 2). The PIWI domain sequence alignment showed that total 15 members namely 7TaAGO1s (TaAGO1a, TaAGO1b, TaAGO1c, TaAGO1d, TaAGO1i, TaAGO1j, TaAGO1k), 4TaAGO5s (TaAGO5c, TaAGO5d, TaAGO5f and TaAGO5g), 2TaAGO7s (TaAGO7a and TaAGO7b), 2TaAGO10s (TaAGO10c and TaAGO10d) possessed the four important active residues (DDH/H) like *Arabidopsis* AGO1(AtAGO1), implying that they might have endonuclease activity[68](Fig. 2). The rest of the 24 TaAGOs showed at least one dissimilarity among these catalytic traid residues. The D760 residue as AtAGO1 in TaAGO8 was changed by Tyrosine (Y) while for others remained same as AtAGO1 (Table 2 and Fig.2). The third residue histidine (H) at 986^th^ position (H986) is missed (“-”) in all TaAGO1e,TaAGO4b,TaAGO6a and TaAGO6e as well as the fourth residue histidine (H) at 798^th^ position(H798) was replace by the proline (P) in TaAGO4b,TaAGO6a and TaAGO6e(Table 2 and Fig.2). The fourth residue in the 798^th^ position (H798) in AtAGO1 of the corresponding position in TaAGO1f, TaAGO1g, and TaAGO1h is replaced by arginine(R) (Table 2 and Fig.2). Unlike AtAGO1, TaAGO2a, and TaAGO2b, TaAGO3 however conserved aspartate (D) as a third catalytic residue instead of histidine (H) in the 986^th^ position (H986) (Table 2 and Fig. 2). On the other hand, the fourth triad residue histidine (H) in the 798^th^ position (H798) is replaced by the amino acid proline (P) in all 11 TaAGO RNAi genes such as 2 TaAGO4s(TaAGO4a and TaAGO4c), 3 TaAGO5s(TaAGO5a TaAGO5b and TaAGO5e), 3 TaAGO6s(TaAGO6b, TaAGO6c and TaAGO6d) and 3 TaAGO9s(TaAGO9a TaAGO9b and TaAGO9c) (Table 2 and Fig.2). Furthermore, tyrosine(Y) was conserved instead of histidine in the 798^th^ position (H798) by TaAGO10a and TaAGO10b proteins (Table 2 and Fig.2). The replacement/mutational changes of the D760, D845, H986 and H798 (DDH/H) catalytic residues in the reported wheat AGO genes are clearly indicating the functional distinction compared to the AGO proteins in Rice and *Arabidopsis*. The further investigation of these genes is demanded for deeper understanding about the PIWI domain nuclease activities as well as the biological functionality in Wheat.

**Table 2:** Comparison of argonaute proteins with missing catalytic residue(s) present in PIWI domain of wheat (*T.aestivum*), rice^b^ and *Arabidopsis*^b^

|  | Wheat |  | Rice |  | *Arabidopsis thaliana* |  |
| --- | --- | --- | --- | --- | --- | --- |
|  | Argonaute | Motif<sup>a</sup> | Argonaute | Motif<sup>b</sup> | Argonaute | Motif<sup>b</sup> |
| 1 | TaAGO1a | DDH/H | OsAGO1 | DDH/P | AGO2 | DDD/H |
| 2 | TaAGO1b | DDH/H | OsAGO2 | DDD/H | AGO3 | DDD/H |
| 3 | TaAGO1c | DDH/H | OsAGO3 | DDD/H | AGO4 | DDH/S |
| 4 | TaAGO1d | DDH/H | OsAGO4a | DDH/P | AGO6 | DDH/P |
| 5 | TaAGO1e | DD-/H | OsAGO4b | DDH/P | AGO9 | DDH/R |
| 6 | TaAGO1f | DDH/R | OsAGO11 | GDH/H |  |  |
| 7 | TaAGO1g | DDH/R | OsAGO13 | -DH/H |  |  |
| 8 | TaAGO1h | DDH/R | OsAGO15 | DDH/P |  |  |
| 9 | TaAGO1i | DDH/H | OsAGO16 | DDH/P |  |  |
| 10 | TaAGO1j | DDH/H | OsAGO17 | HDR/C |  |  |
| 11 | TaAGO1k | DDH/H | OsAGO18 | DDH/S |  |  |
| 12 | TaAGO2a | DDD/H |  |  |  |  |
| 13 | TaAGO2b | DDD/H |  |  |  |  |
| 14 | TaAGO3 | DDD/H |  |  |  |  |
| 15 | TaAGO4a | DDH/P |  |  |  |  |
| 16 | TaAGO4b | DD-/P |  |  |  |  |
| 17 | TaAGO4c | DDH/P |  |  |  |  |
| 18 | TaAGO5a | DDH/P |  |  |  |  |
| 19 | TaAGO5b | DDH/P |  |  |  |  |
| 20 | TaAGO5c | DDH/H |  |  |  |  |
| 21 | TaAGO5d | DDH/H |  |  |  |  |
| 22 | TaAGO5e | DDH/P |  |  |  |  |
| 23 | TaAGO5f | DDH/H |  |  |  |  |
| 24 | TaAGO5g | DDH/H |  |  |  |  |
| 25 | TaAGO6a | DD-/P |  |  |  |  |
| 26 | TaAGO6b | DDH/P |  |  |  |  |
| 27 | TaAGO6c | DDH/P |  |  |  |  |
| 28 | TaAGO6d | DDH/P |  |  |  |  |
| 29 | TaAGO6e | DD-/P |  |  |  |  |
| 30 | TaAGO7a | DDH/H |  |  |  |  |
| 31 | TaAGO7b | DDH/H |  |  |  |  |
| 32 | TaAGO8 | YDH/P |  |  |  |  |
| 33 | TaAGO9a | DDH/P |  |  |  |  |
| 34 | TaAGO9b | DDH/P |  |  |  |  |
| 35 | TaAGO9c | DDH/P |  |  |  |  |
| 36 | TaAGO10a | DDH/Y |  |  |  |  |
| 37 | TaAGO10b | DDH/Y |  |  |  |  |
| 38 | TaAGO10c | DDH/H |  |  |  |  |
| 39 | TaAGO10d | DDH/H |  |  |  |  |

<sup>a</sup>Motifs are correspond to conserved D760, D845, H986/H798 of Arabidopsis AGO1; D(aspartate), H(Histidine), K(Lysine), A(Alanine), P(Proline), Q(Glutamine), G(Glycine), R(Arginine), C(Cysteine), S(Serine); ‘-’ represents the missing catalytic residue
<sup>b</sup> Reviewed in Kapoor *et al* (2008)

Finally, total 16 TaRDR proteins however possessed an RdRp domain (Table 3). The gene length of the TaRDR proteins varied from 15893bp for TaRDR2b to 1707 bp for TaRDR1d with their corresponding potentially encoding 1127 and 484 amino acids respectively (Table 1). Additionally, TaRDR proteins were studied to identify the presence of the catalytic motif in the RdRp domain. The sequence alignment revealed that all members of TaRDR1, TaRDR2, and TaRDR6 groups shared a common DLDGD motif in the catalytic domain, whereas the proteins TaRDR3, TaRDR4, and TaRDR5 retained DFDGD motif (Fig. 3).

**Table 3.**
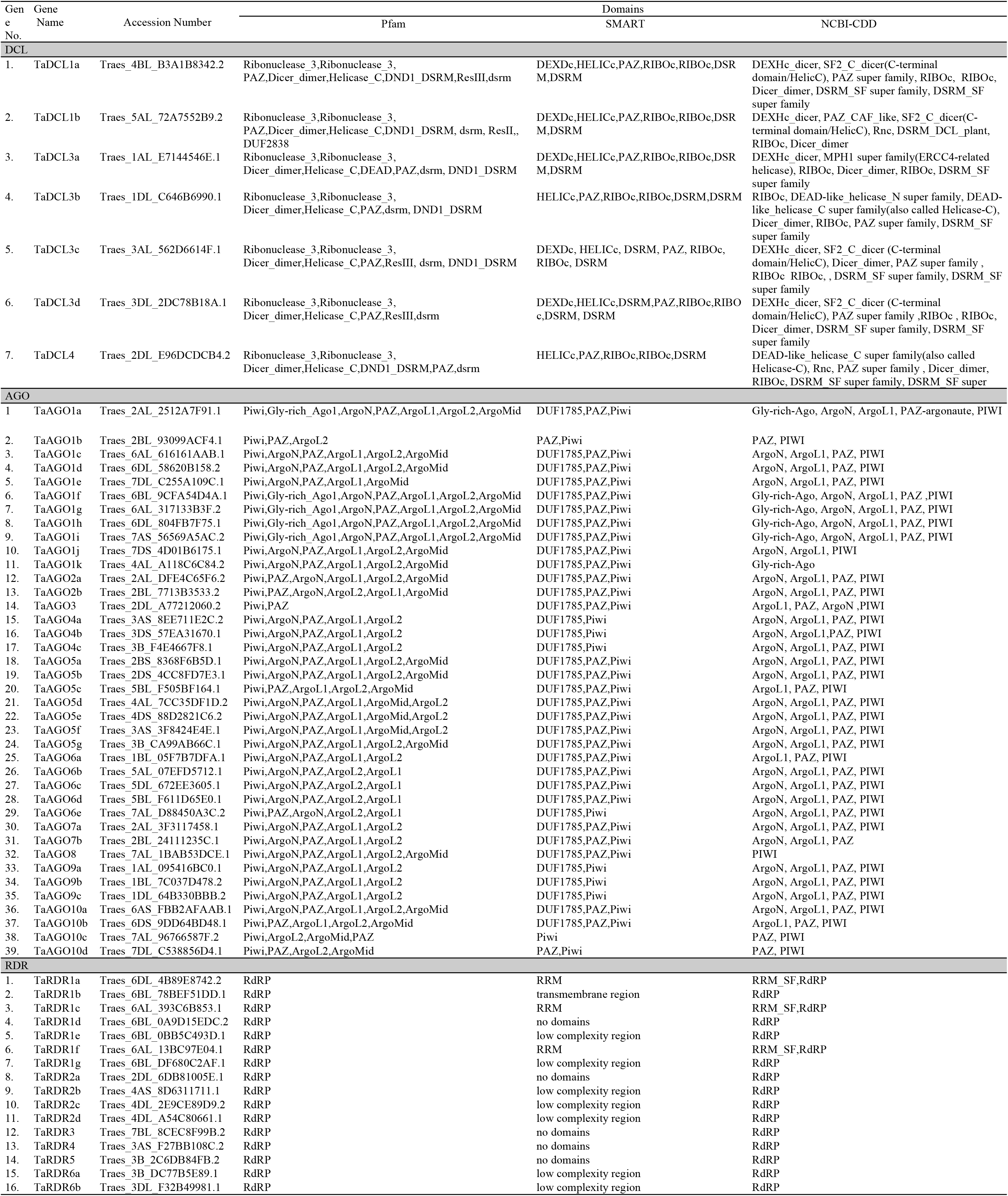
List of predicted conserved functional domains of the DCL, AGO and RDR proteins in wheat(*T.aestivum*) using Pfam, SMART and NCBI-CDD.

**Fig.3.**
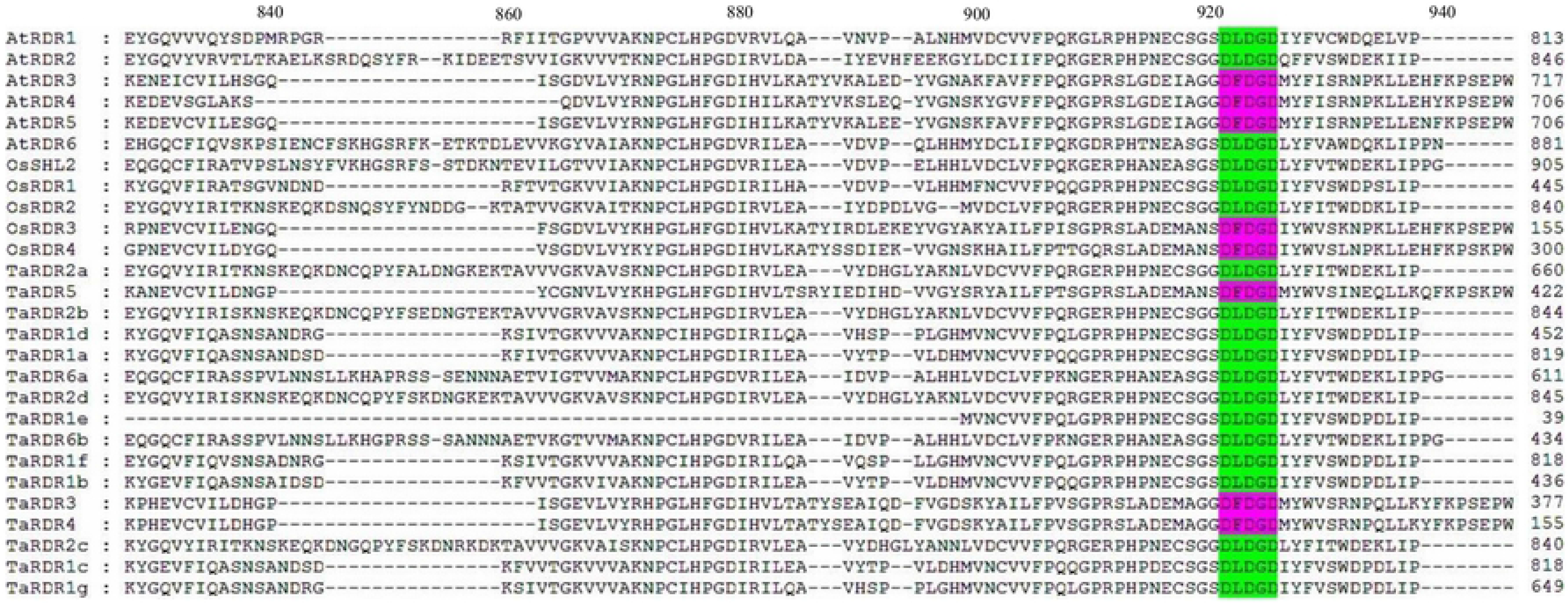
Alignment of catalytic regions in RdRp domains of RDR proteins of *T. aestivum*, rice, and *A. thaliana*. The conserved motif DLDGD and DFDGD were colored. The alignments were performed using MEGA 7.0. A “−” indicates of an amino acid missing of the corresponding protein sequence. The locations analogous to each protein are mentioned at the end of line.

### 3.2 Phylogenetic tree analysis of the putative RNAi genes in *T.aestivum*, *A.thaliana* and rice

In order to infer the evolutionary history of the three groups of RNAi genes in wheat (*T. aestivum*) with those of the *Arabidopsis* and rice homologs, three independent phylogenetic trees were created from the multiple sequence alignment. Encoded protein sequences (S1, S2 and S3 data) of all RNAi genes were used to construct the trees using neighbor-joining(NJ) method with 1000 bootstrap replications (Fig. 4).The TaDCL proteins showed high sequence conservation compared with their *Arabidopsis* and rice counterpart. Fig. 4A shows that the 7TaDCLs were obviously classified into four distinct clades. All the TaDCL members of the corresponding group are clustered with the similar ortholog AtDCLs and OsDCLs at a high similarity context. In relation to the phylogenetic association and sequence homology with AtDCLs, the TaDCLs were named as TaDCL1a, TaDCL1b, TaDCL3a, TaDCL3b, TaDCL3c, TaDCL3d, and TaDCL4. However, there were no TaDCL2 gene(s) corresponding to AtDCL2 and OsDCL2.

**Fig.4.**
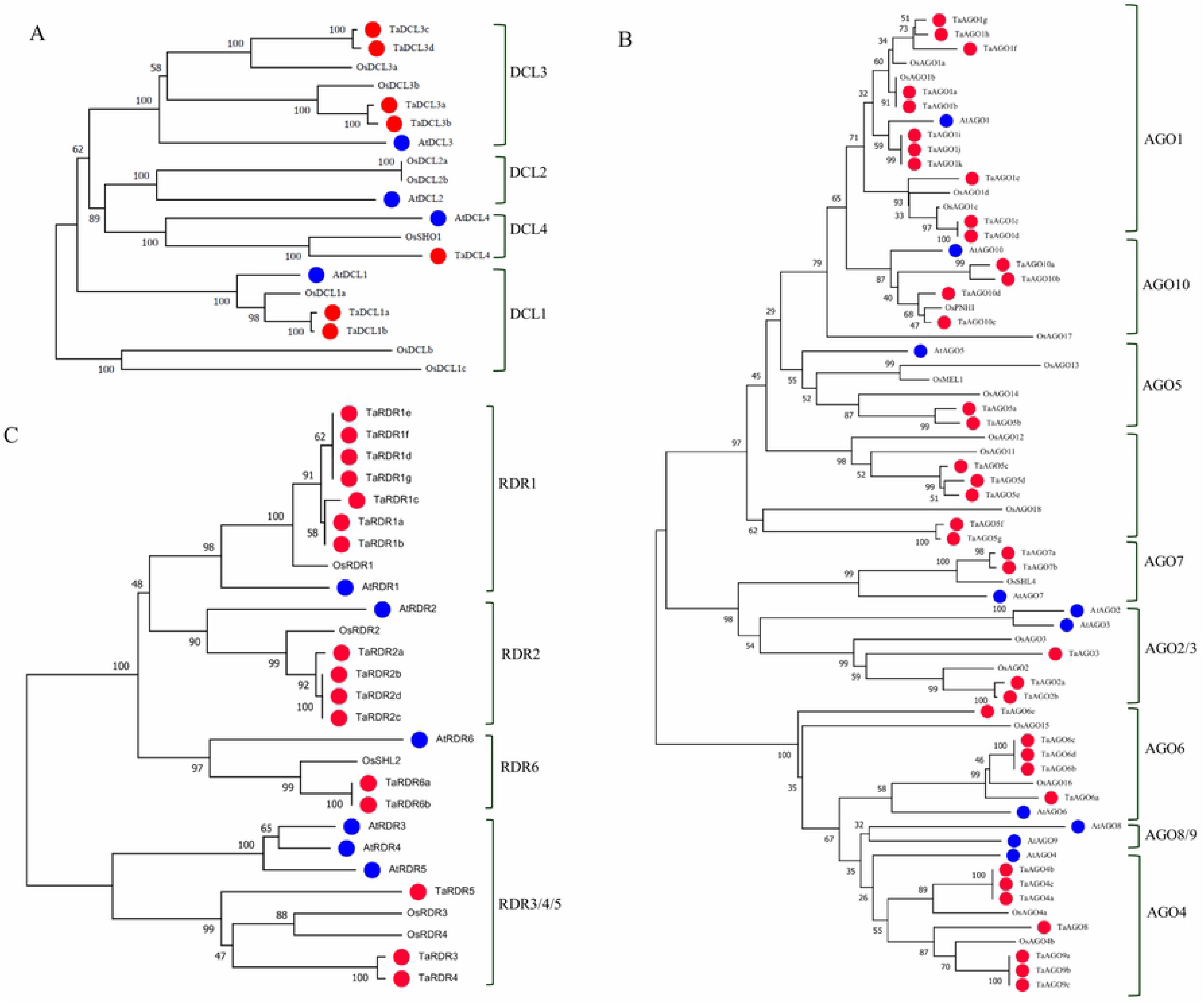
Phylogenetic trees of all *A. thaliana*, rice (*Oryza sativa*) wheat (*T. aestivum*) RNAi gene families. (A) DCL (B) AGO and (C) RDR RNAi group. The trees were produced by MEGA 7.0 software using Neighbor-Joining (NJ) method with bootstrap of 1000.*T. aestivum* proteins are indicated with a red colour circle before the corresponding taxon names.

Based on NJ phylogenetic analysis and the protein sequence homology with AtAGOs, the 39 candidates of TaAGO family consisted of 11 AGO1s (TaAGO1a-TaAGO1k), 2AGO2s (TaAGO2a and TaAGO2b), one AGO3 (TaAGO3), 3AGO4s (TaAGO4a, TaAGO4b and TaAGO4c), 7 AGO5s (TaAGO5a-TaAGO5g), 5AGO6s (TaAGOa-TaAGO6e), 2AGO7s (TaAGO7a and TaAGO7b), one AGO8 (TaAGO8), 3AGO9s (TaAGO9a-TaAGO9c) and 4AGO10s (TaAGO10a-TaAGO10d) for 8 clusters (Fig. 2B). It is however observed from the phylogenetic tree that TaAGO8 and 3TaAGO9s (TaAGO9a-TaAGO9c) form a group with the nearest taxa OsAGO4b not with the corresponding AtAGO8 and AtAGO9. On the other hand, it is unusually obvious that TaAGO5a and TaAGO5b group with OsAGO14. TaAGO5c, TaAGO5d, TaAGO5e form a group with OsAGO11 and OsAGO12 as well as TaAGO5f and TaAGO5g specifically with taxa OsAGO18 but not with any AtAGOs. The characteristic of these genes may however indicates that TaAGO8, 3TaAGO9s, and 7TaAGO5s are the monocot specific that have originated during evolutionary process[7,10].

Finally, like DCL and AGO, RDR genes in *T. aestivum* were named after the *Arabidopsis* homologs, which were the evolutionary neighboring in the phylogenetic tree, and exhibit the top protein sequence homologies. Fig. 2C illustrates that the phylogenetic tree however produced from the RDR protein sequences is separated into four major clusters. RDR1 clade has seven members (TaRDR1a, TaRDR1b, TaRDR1c, TaRDR1d, TaRDR1e, TaRDR1f, TaRDR1g,) form a cluster with their AtRDR1 and OsRDR1 counterparts. Clade RDR2 has four members (TaRDR2a, TaRDR2b, TaRDR2c, TaRDR2d,) and clearly generates a distinct cluster with their AtRDR2 and OsRDR2 homologs. TaRDR3, TaRDR4, and TaRDR5 together form a cluster with their similar genes to AtRDR3, AtRDR4, AtRDR5, OsRDR3, and OsRDR4. Clade RDR6 has two members TaRDR6a and TaRDR6b produced a distinct group with AtRDR6 and OsSHL2 genes in the phylogenetic tree imply that these TaRDR protein families covey the similar genetic information of their AtRDR and OsRDR counterparts.

### 3.3. Discovery of conserved domains and motifs of the putative DCL, AGO and RDR proteins in *T.aestivum*

Protein domains have functional importance in protein structure. They are made up of a chain of amino acids of independent molecular folding. Domains have unique function in respective protein. Functional conserved domain analysis for protein structure in *T.aestivum* of the predicted TaDCLs, TaAGOs and TaRDRs were performed using Pfam, SMART and NCBI-CDD databases. The obtained search results are listed in Table 3.

It is observed from the search results from all three databases that all TaDCL protein families possessed all most all assumed six common domains viz., DEAD box/ResIII, Helicase-C, Dicer-dimer, PAZ, RNase III and DSRM (Table 3 and Fig.5). These were also conserved by the 4 DCL protein families in model plant *Arabidopsis*[7,10,70]. Previous study suggests that these functional domains are known as the highly important plant DCL domains for protein structure[7,10,11,70]. Ribonuclease-3(RNaseIII) domain characteristics of TaDCL proteins might enable them to cleave dsRNA to generate small interfering RNAs(siRNAs) that play crucial functions to regulate gene expression in plants[10,11,64]. On the other hand, earlier investigations revealed that Argonaute proteins usually retain a Piwi-Argonaute-Zwille (PAZ) domain and a PIWI domain[71,72]. Functional domain search results (Table 3) and domain organization by Pfam (Fig. 5) showed that all 39 TaAGO protein families also hold the two most common domains, viz., N-terminus PAZ and C-terminus PIWI domain similar to that of *Arabidopsis* and rice known as potential characteristic of plant AGO families[7,13]. Additionally, a DUF1785 domain also can be recognized before PAZ domain in all TaAGO proteins except for TaAGO1b, TaAGO10c and TaAGO10d encodes the shortest amino acids chain, which may however be for the possible of loss of its N-terminal sequence during evolution

**Fig.5.**
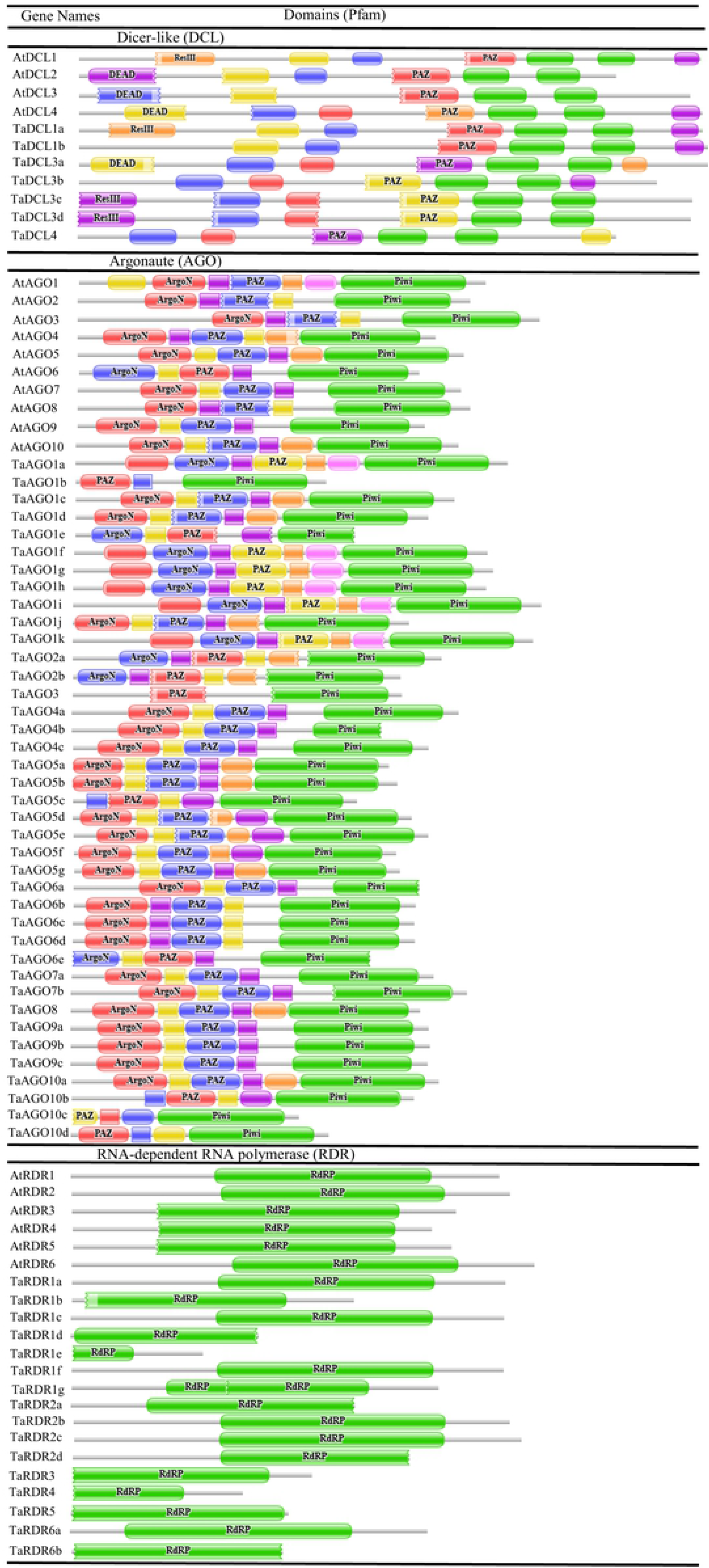
Domain organization of DCL, AGO and RDR protein families in wheat (*T.aestivum*). Conserved functional domains were predicted from Pfam database. Domains are indicated in different color boxes with their corresponding name inside the box.

Besides, the *T.aestivum* genome encrypted 16 TaRDR protein families sharing a mutual motif analogous to the catalytic β’ subunit of RNA-dependent RNA-polymerases(RdRp)(Table 3 and Fig.5), which is also supported by the earlier study[33].

Motifs in a DNA, RNA or a protein sequence encode different biological functions[73]. Their identification and characterization is significant to study molecular interactions in cell such as the regulation of gene expression. These motifs are also known as the small active site of an enzyme or a structural component essential for accurate folding of the protein and molecular evolution[73]. In this work, Multiple EM for Motif Elicitation (MEME) was used to discover motifs for each group of TaDCL, TaAGO and TaRDR protein sequences along with that of RNAi protein families of *Arabidopsis*. The search was done for each member for each of the family and 20 motifs were predicted for each of the proteins. It is observed that most of the motifs were conserved in each of the TaDCL protein members and distributed in order in almost all four subfamilies of the DCL protein family like *Arabidopsis* (Fig. 6). Wheat DCL4 protein subfamily, TaDCL4, however was identified to possess a few C-terminal motifs reorganized in comparison of the paralogs AtDCL4, which was also supported by the previous demonstration[10]. All TaDCL protein subfamilies retain 20 conserved motifs with the exception of that TaDCL3b and TaDCL4 contain 19 and 16 motifs respectively (Fig.6).

**Fig.6.**
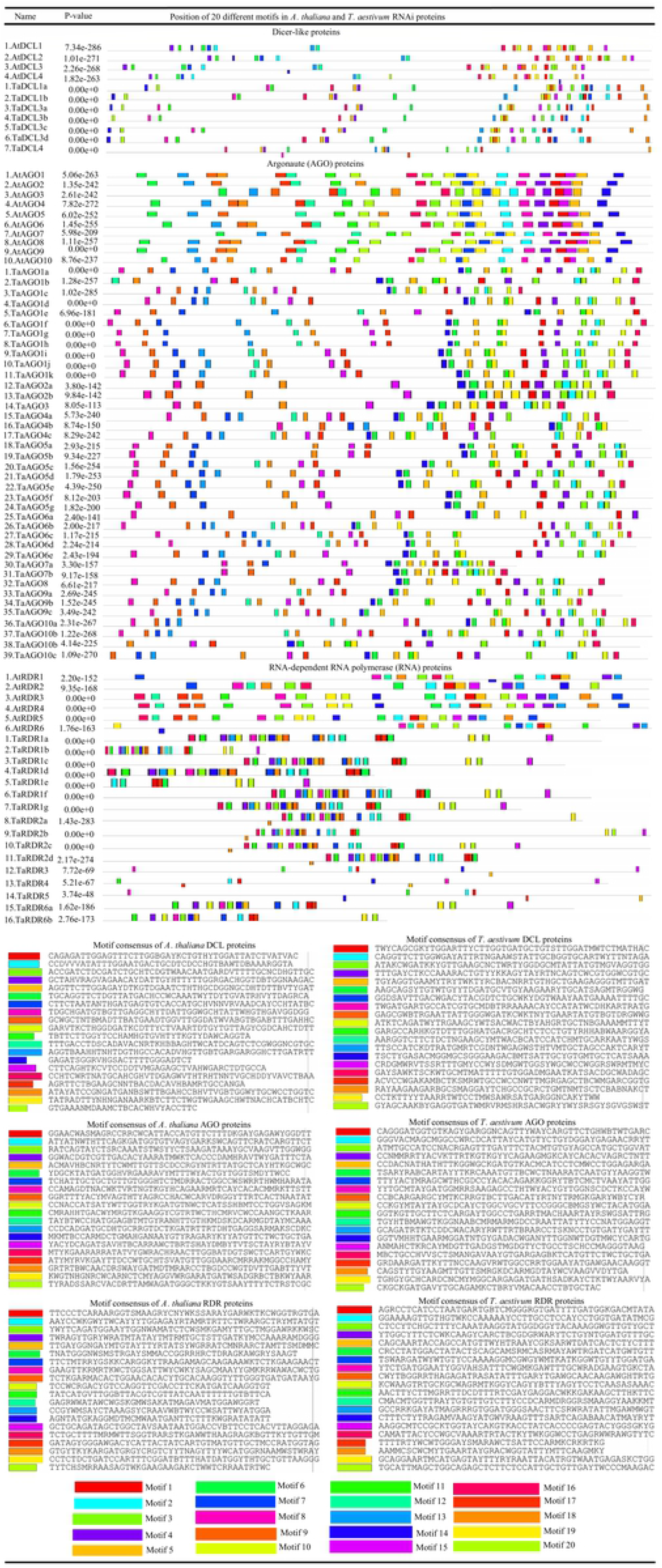
Discovery of 20 conserved motif structures. Motifs with consensus amino acid sequence in wheat DCL, AGO and RDR protein families predicted by the web-based MEME suite. Each motif in the protein domain is represented by the color box and the order of the motifs corresponds to position of motifs in individual protein sequence.

On the other hand, out of 39 TaAGO protein subfamilies, the maximum 28 protein members of wheat DCL, AGO and RDR retained 16-20 conserved motifs in their protein domain structures with some variations in other 11 subfamilies. High conservation was found in TaAGO1, TaAGO9, and TaAGO10 family members with some exceptions that TaAGO1b, TaAGO1e, and TaAGO10c contained 16, 14, and 15 motifs respectively (Fig.6).

Motif search using MEME for the RDR protein family in wheat demonstrates that 10 TaRDR subfamilies possess 17-20 functional motifs in the RdRp domain similar to that of the AtRDR protein members (Fig.6) but with little high distinction that TaRDR1e, TaRDR3, TaRDR4, and TaRDR5 contain only 7-8 motifs. The study of sequence motifs of the protein subfamilies of each group of RNAi genes, it suggests that there is a tendency of high conservation of functional motifs in respective domains within the members of same subfamilies in TaDCL, TaAGO and TaRDR groups that are assumed to hold and perform same structural functions. Therefore, discovery and broad study of these motifs is important to structure models of cellular processes at the molecular scale and to identify the mechanisms of diseases in eukaytic groups[58,73].

### 3.4. Assembly of exon-intron structure of T. aestivum DCL, AGO and RDR Genes

The exon-intron organization of TaDCL, TaAGO and TaRDR genes in wheat (*T.aestivum*) was studied using GSDS2.0 web-based tool for more broad understanding regarding their probable structural progression and level of similarity with the respective RNAi genes of *Arabidopsis thaliana*. Our analysis for all three gene families illustrated that intron and exon numbers were usually conserved among the members of the same group though varied considerably in different groups of the same family that is similar to those of the AtDCL, AtAGO, and AtRDR gene families. The intron numbers varied from 9 to 18 among TaDCL protein members while 4 AtDCLs possessed 11-17 introns (Table 1 and Fig. 7). The two members of TaDCL1 sub-group contained 9 introns, which is the lowest in the group whereas only TaDCL3a contained the double introns compared to this sub-group. Other four members retained 15/16 introns (Table 1 and Fig. 7). On the other hand, almost all TaAGO protein families also follow the same range of introns of their AtAGOs counterpart commonly measuring 13-18 although TaAGO2a/2b,TaAG03 and TaAGO7a/7b displayed only lowest intron numbers among all argonaute gene members in wheat(*T.aestivum*) like AtAGO2, AtAGO3 and AtAGO7(Table 1 and Fig.7). The members of the TaAGO1 sub-family maintained high similarity conservation in containing introns16-18 as in AtAGO1 with exception 11 introns in TaAGO1b (Table 1 and Fig. 7).Furthermore all members of the rest of the sub-families follow almost the same pattern in containing introns and exons of their *Arabidopsis* counterpart. Among the TaRDR protein elements, TaRDR3 contained the highest number of 10 introns followed by 9 and 5 introns in TaRDR5 and TaRDR4 respectively (Table 1 and Fig.7). Interestingly, all gene members of TaRDR1 and TaRDR6 sub-group had no introns though AtRDR1 possessed 2 introns and AtRDR6 had 0 introns that might be of some significant implication that these gene sub-families may be responsible for any organ or tissue specific expression characteristics and bear any particular disease resistance potentiality (Table 1 and Fig.7). Earlier investigation suggested CaRDR1 possessed important role in pepper resistance against TMV[74].It was however observed that the exon-intron structure of TaDCL, TaAGO and TaRDR genes was very similar within members of the same groups though substantially different through various groups of the same family indicate that these gene families particularly TaAGO and TaRDR families have experienced recurrent gene replication and recombination during evolution.

**Fig.7.**
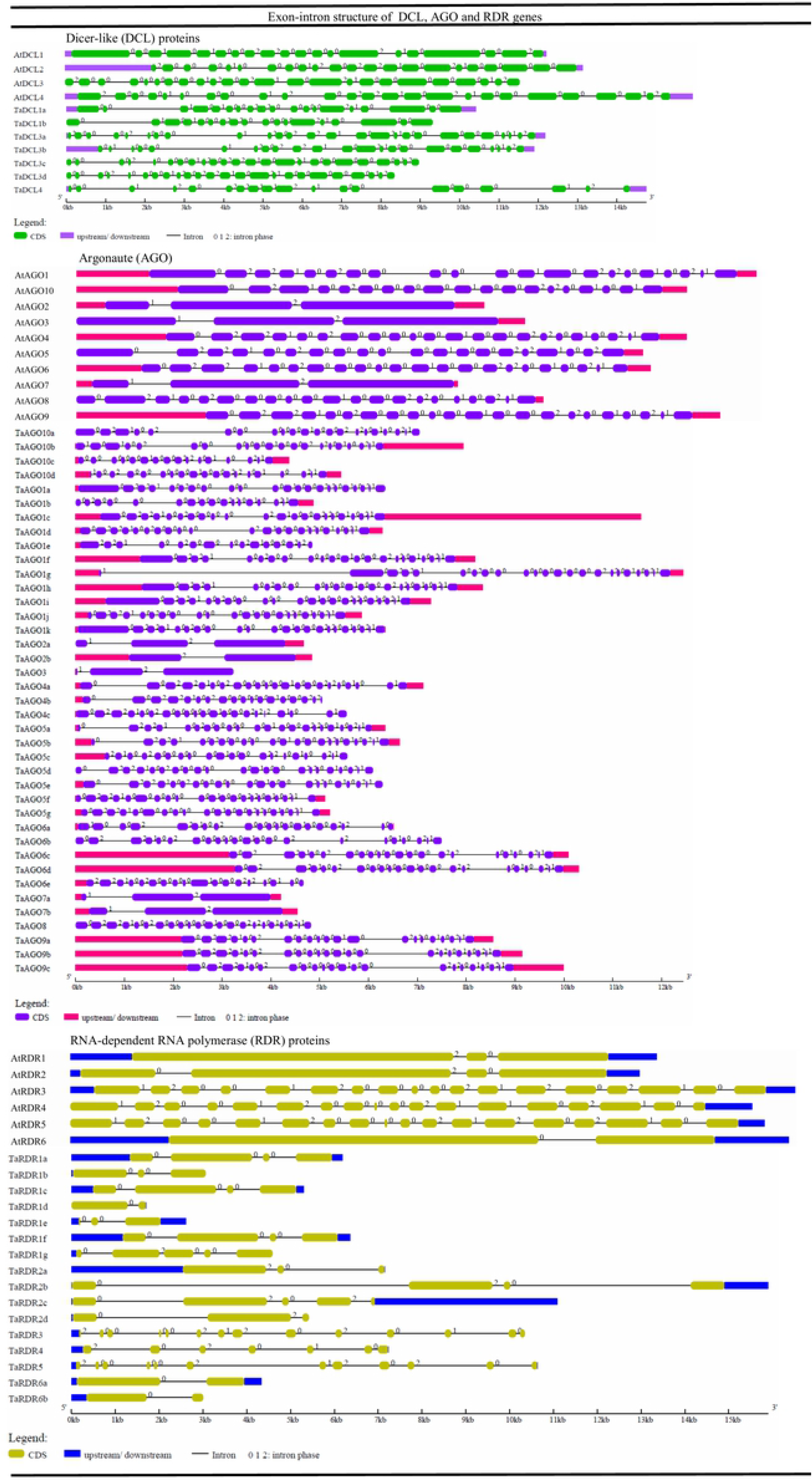
Exon-intron structure of *T. aestivum* RNAi genes corresponding to *A. thaliana* DCL, AGO and RDR genes. Exons (green box), intron (black lines), and intron phases (0, 1, and 2) are mentioned. The gene structure was estimated using the online GSDS1.0 server by considering their full-length genomic/DNA and coding sequences (CDS).

### 3.5. Gene set enrichment analysis of the RNAi genes in wheat (*T.aestivum*)

Gene set enrichment analysis(GESA) such as gene ontology (GO) analysis was conducted to demonstrate that how the predicted RNAi genes in *T.aestivum* are involved in different biological processes (BP), what molecular functions(MF) they perform with their interrelations and what are the cellular components where these molecular functions occur(S4 data, S1 Fig. and Fig.8). An interconnected relationship using QuickGO web-based tool from less specific concepts to more specific concepts among the GO terms and the corresponding gene ontologies are demonstrated using the three hierarchical GO slim trees for BP (S1A Fig.), MF (S1B Fig.), and CC (S1C Fig.). It is shown from the GO analysis that 11 gens are involved in post-transcriptional gene silencing (PTGS) pathway (GO:0016441;p-value: 5.00e−24), 9 genes are responsible for RNAi interference mechanism(GO:0016246;p-value:3.60e−22) and 11 genes are linked to gene silencing pathway(GO:0016458;p-value:1.30e−19)(S4 data). Post-transcriptional gene silencing (PTGS) in plants is an RNA-degradation technique that displays connections to RNA interference (RNAi) [75,76]. GO slim hierarchy tree illustrates that RNA interference is a sub-type of PTGS ultimately connected to gene expression (GO: 0010467; p-value: 7.20e−05) (S1A Fig.). There are also a number of biological processes are related to be occurred by the predicted RNAi genes in wheat (*T.aestvum*) some of which important for wheat plant are response to stimulus (GO: 0050896; 0.00017), immune system process (GO: 0002376; 3.70e−14) and developmental process (GO: 0032502, 5.40e−06) (S4 data and S1A Fig).GO tree displayed that response to stimulus is termed into three important sub-types GO such as response to stress(GO:0006950; 2.80e−05) linked to 10 RNAi genes, response to external stimulus(GO:0009605; 7.30e−11) and response to biotic stimulus(GO:0009607; 1.20E−11). These three terms related to defense response to virus (GO: 0051607; 4.80e−22) called ultimately virus induced gene silencing (GO: 0009616; 6.10e−20). It is also obvious from the GO tree that response to stimulus and immune system process is associated to the defense response to other organism (GO: 0098542; 3.90E−13) (S1A Fig.). GO analysis for BP also suggests that some very important functions are maintained by the RNAi genes in wheat (*T.aestivum*), viz., post-embryonic development (GO: 0009791; 6.90E−05), anatomical structure development (GO: 0048856; 0.00014) which are eventually connected to wheat plant developmental process (GO: 0032502; 2.87e−04) (S1A Fig.) Previous investigation also reported that *Arabidopsis* DCL and RDR RNAi genes are actively associated to defense response to various biotic, abiotic stressors and play role for developmental progression[41,42,50–54].

**Fig.8.**
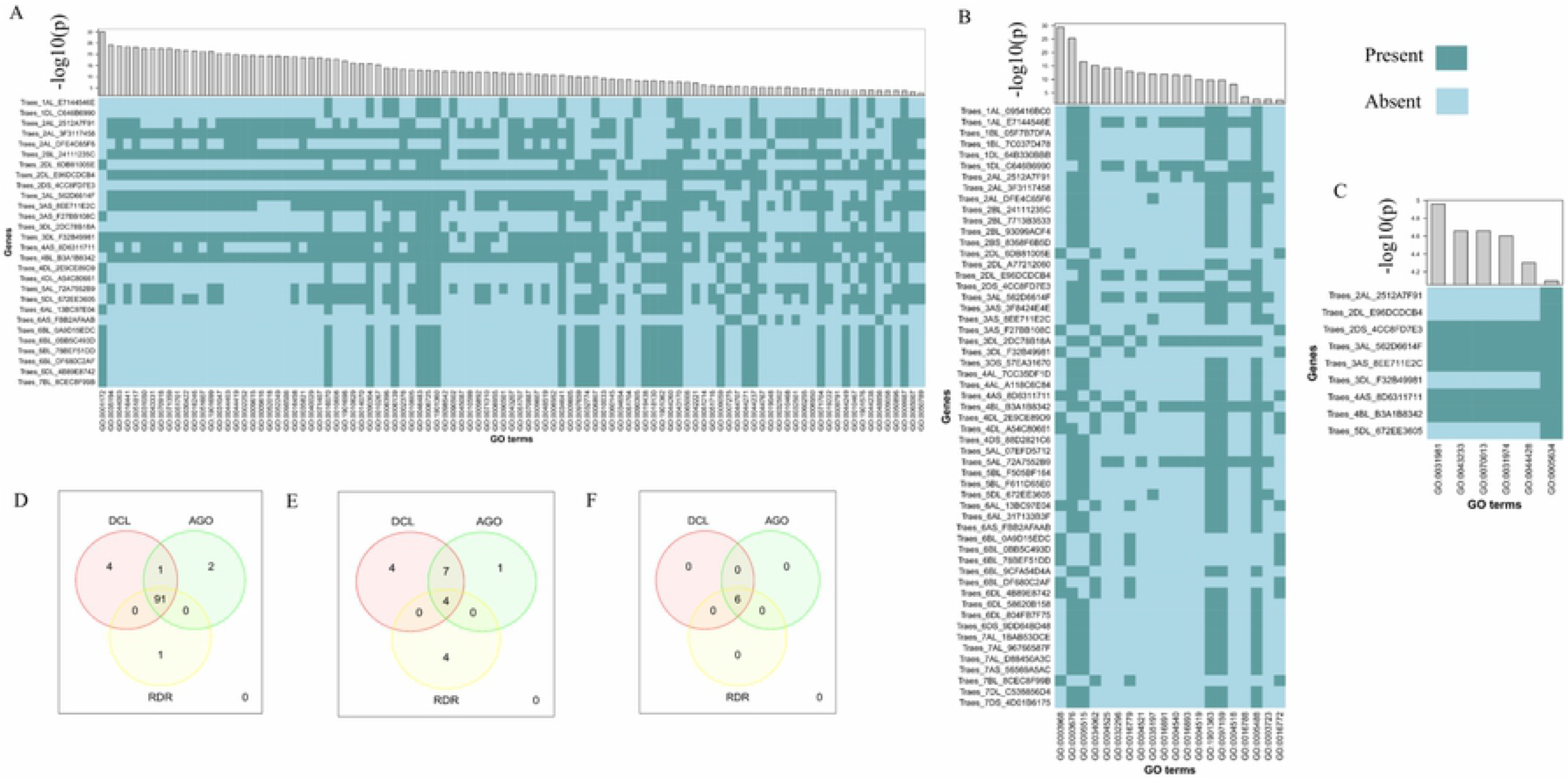
Heatmap and Venn diagram for the predicted GO terms corresponding to the wheat RNAi genes. (A) biological process (B) molecular function (C) cellular components whether the genes are related (Present) or not (Absent). Histograms corresponding to each GO terms represent the p-value (−log (p-value)) showed in the opposite site of GO terms in the heatmap. The Venn diagrams display the common GO terms shared by three gene families in terms of (D) biological process (E) molecular functions (F) cell components.

GO enrichment analysis showed that 8 putative RNAi genes out of 62 are associated to perform endonuclease type molecular function (GO:0004519;p-value: 1.20e−10)(S4 Data) that have positive link to the RNA-induced silencing complex(RISC-mediated) mediated cleavage actions into cell[7,10,76,77].This multi-protein molecular compartment play significant role in mRNA degradation through that regulate gene expression. Argonaute proteins also act as attaching themselves with siRNAs in RISC for cleavage defined as endonucleolytic function that results in completion of PTGS for target mRNAs substrate[78]. GO analysis also demonstrates that three essential molecular functions(MF), viz., nucleic acid binding(GO:0003676; 5.00e−26), protein binding(GO:0005515; 2.80e−17), RNA binding(GO:0003723;p-value:0.00257) are associated to 44, 42 and 7 predicted RNAi genes in wheat(*T.aestivum*)(S4 Data) which implied that RNAi proteins are actively engaged in RISC as well as gene silencing. Nucleic acid binding is also of two types: heterocyclic compound binding (GO: 1901363; 1.80e−10) and organic cyclic compound binding (GO: 0097159; 1.80e−10) (S4 Data). GO ancestry tree showed that all these MFs are the child terms of the GO term binding (GO: 0005488; 0.00217) (S1B Fig.).There is another MF termed as ribonuclease III activity (GO: 0004525; p-value: 7.00E−15) is related to 7 RNAi genes that is the child GO term of double-stranded RNA-specific ribonuclease activity (GO: 0032296; p-value: 7.00e−15) and nuclease activity (GO: 0004518; p-value: 6.70e−09) (S1B Fig.). The identified TaAGO proteins possess the two common domains like other plant AGO proteins such as PAZ and PIWI that perform essential functions to build protein complex with RNA or DNA template. The PAZ domain has a particular nucleic acid-binding fold which help domain to bind to the certain site of the nucleic acids[79,80]. Different molecular functions are usually tend to occur in various cellular components (CCs). GO analysis also suggests that the different MFs are predicted to happen significantly in nucleus (GO: 0005634; 8.00e−05), intracellular organelle lumen (GO: 0070013; 2.20e−05), organelle lumen (GO: 0043233; 2.20e−05), and membrane-enclosed lumen (GO: 0031974; 2.50e−05) (S1C Fig.).

The Venn diagram also illustrates that TaDCLs, TaAGOs and TaRDRs shared significant number of GO pathways. For BP, there were 91 GO enrichment pathways in general (Fig. 7D) shared by the identified RNAi genes in wheat (*T.aestivum*), that suggests that RNAi genes are significantly associated to many important BPs. Furthermore, it were shown that the identified putative RNAi genes displayed a pool of mutual pathways for performing MFs and these MFs area predicted to occur in 6 common CCs(Fig. 8E and Fig. 8F). GESA however provided insights about the identification of the involvement of 62 wheat (*T.aestivum*) RNAi genes in different biological processes; engagement in performing a number of molecular actions in different cellular components for the survival of wheat plant in diverse environments.

### 3.6 Prediction of sub-cellular location of the TaDCL, TaAGO and TaRDR gene families in wheat (*T.aestivum*)

Prediction of sub-cellular localization (SCL) helps to know the presence or absent status of the identified RNAi genes in certain cellular organs in plants. A number of RNA silencing machinery genes in wheat (*T.aestivum*) were found locating in more than one cellular locations (Fig. 9). It is however observed that most of the RNA silencing genes were found to be located in cytosol (DCL 71.4 %, AGO 87.2 % and RDR 87.5 %) followed by plastid (DCL 14.3 %, AGO 33.3 % and RDR 31.2 %). A few of the AGO (20.5 %) and RDR (12.5 %) genes were located in mitochondria but no DCL protein was found in mitochondria. On the other hand, ER, golgi and peroxisome contain no RNA silencing genes (**Fig. 9D and S2 Table**). Cytosol is the place of maximum metabolism in plants and most of the proteins in the cell are located in cytosol[81,82]. Since the PTGS takes place in the cytoplasmic region [77], this suggests that the identified putative RNAi genes are actively engaged in PTGS process.

**Fig.9.**
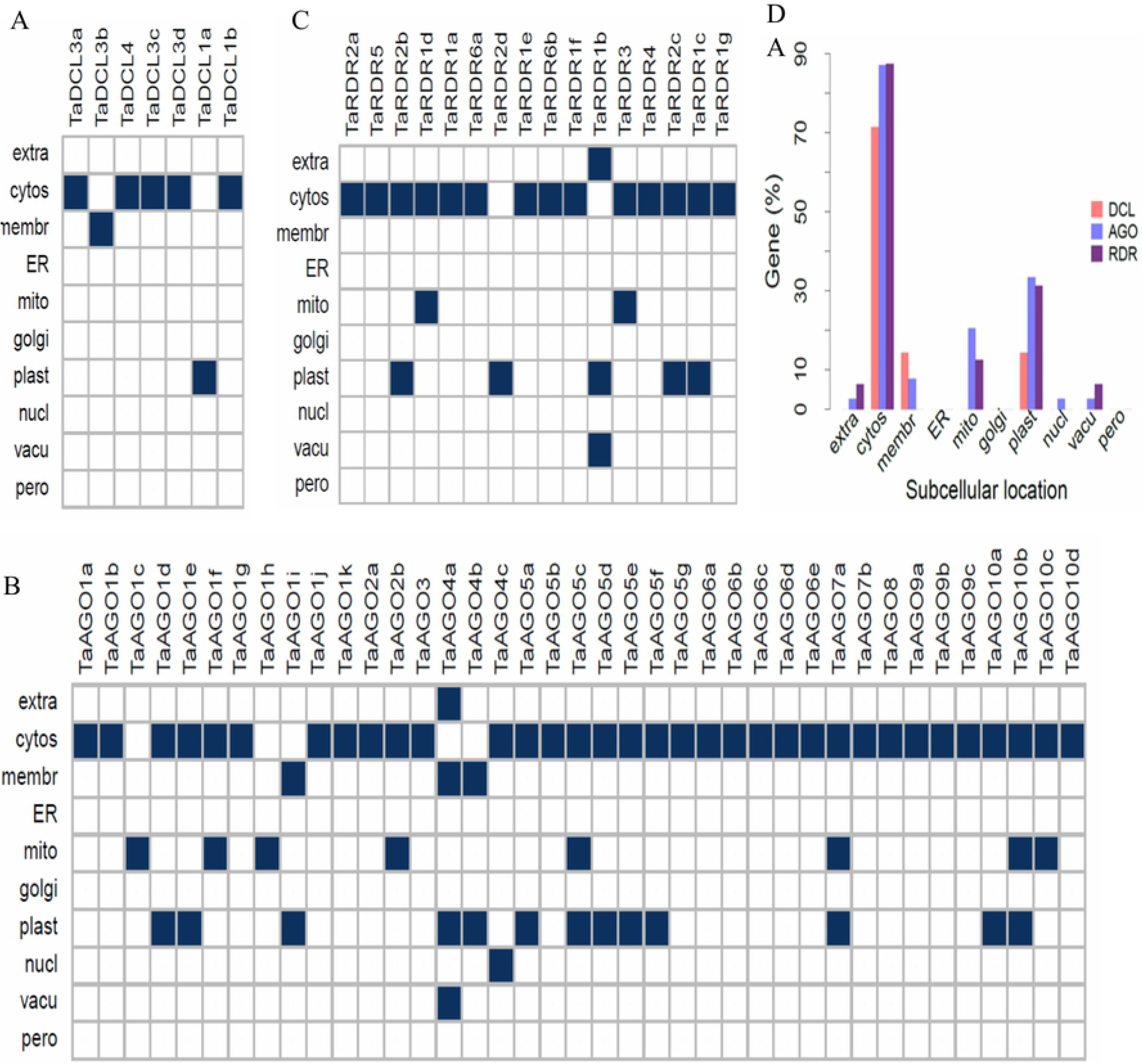
Prediction of subcellular locations of TaDCL, TaAGO and TaRDR gene families. Prediction were made for cytosol (cytos), endoplasmic reticulum (ER), extracellular (extra), golgi apparatus (golgi), membrane (membr), mitochondria (mito), nuclear (nucl), peroxisome (pero), plastid (plast) and vacuole (vacu) for each of the (A) DCL, (B) AGO and (C) RDR gene families in *T. aestivum* with the help of PSI and R-3.6.3.

### 3.7 Exploration of gene regulatory relationship between TFs and RNAi genes in wheat (*T.aestivum*)

A transcription factor (TF) acts as a key for a particular gene expression in lives. It is also known as a DNA-binding amino acid chain, which particularly works attaching with a particular *cis*-acting component in the promoter site of a gene for gene expression in plants and other eukaryotic groups[83]. Recent research in plant molecular biology has attracted genomics researchers to study the important transcription factor families to study the promoting and controlling power of gene expression against different biotic and abiotic stresses such as fungi, bacteria, viruses, drought, salinity, cold, hormone, pathogenesis, cell growth, and development in higher plants. Therefore, identification and characterization of TF families regulating the RNAi genes is however essential to study the gene silencing process in wheat (*T.aestivum*).

In this analysis, 375 TFs were identified those regulate the predicted RNAi genes in wheat (*T.aestivum*) (S5 Data and S2 Fig.) for particular gene expression. The identified TFs were classified into 27 groups based on the TF families (S2 Fig.). Among the putative TF families, ERF, MIKC-MADS, C2H2, BBR-BPC, MYB, Dof, LBD,CPP and AP2 families are assumed highly contribute in regulating RNAi genes for specific gene expression in wheat (*T. aestivum*) in diverse conditions as these families bind with higher number of RNAi genes in wheat. Out of these top 7 TF families, first three one, ERF possesses 157(41%), MIKC-MADS possesses 38(10%) and C2H2 retains 34(9%) TFs, which are in leading position accounted for nearly 61% of the total identified TFs (S3 Table). This, implied that those set of TF families could be high potential in regulating gene expression in terms of predicted RNAi genes in wheat (*T.aestivum*).

Furthermore, a regulatory gene network was created using Cytoscape to analyze in-depth relationship among the TFs and RNAi genes in wheat(*T.aestivum*).This suggests that there is some diverse relationships among the predicted groups of TFs and predicted RNAi genes (Fig. 10) in wheat. The ERF, top most TF family, largely binds to the genes TaDCL3a (18 TFs), TaAGO7a (17 TFs), TaDCL1a (15 TFs), TaAGO9b (15 TFs), TaRDR2b (16 TFs), TaAGO2a (13 TFs), and TaAGO10b with 13 TFs (Fig.10 and S2 Fig.). MIKC-MADS, second top most TF family, also binds with the higher number of TaRNAi genes among those TaAGO1f(6 TFs), TaAGO6c(3 TFs), TaAGO6d(3 TFs), TaRDR1e(4 TFs) and TaRDR3(3 TFs) followed by C2H2, BBR-BPC and MYC. It is also observable that other than these leading TF families in wheat, the two families, viz., AP2 (one TF with each TaDCL3a, TaAGO1f, TaAGO6b, TaAGO6d, TaAGO6e TaAGO9c, TaAGO10c, TaRDR1f, TaRDR2d and TaRDR3) and bZIP (4 TFs with TaAGO2b) are predicted to bind with some specific *cis*-acting elements in the RNAi genes to response some physiological and biochemical stimuli in wheat for regulating gene expression (S2 Fig.). Some previous study suggested that a few TFs regulate the expression of a number of important genes for some particular physiological and biochemical reasons[83,84]. Earlier investigation also reported that both AP2 and bZIP TF families contribute in gene regulation for development and resistance response to various environmental stressors[83]. From the analysis results, it is however observed that AP2 TF family only interacts with TaAGO2b (S2 Fig.) in wheat that might be predicted that the DNA-binding domain of this TF family is responsible to play for regulating gene in a specific situation.

**Fig.10.**
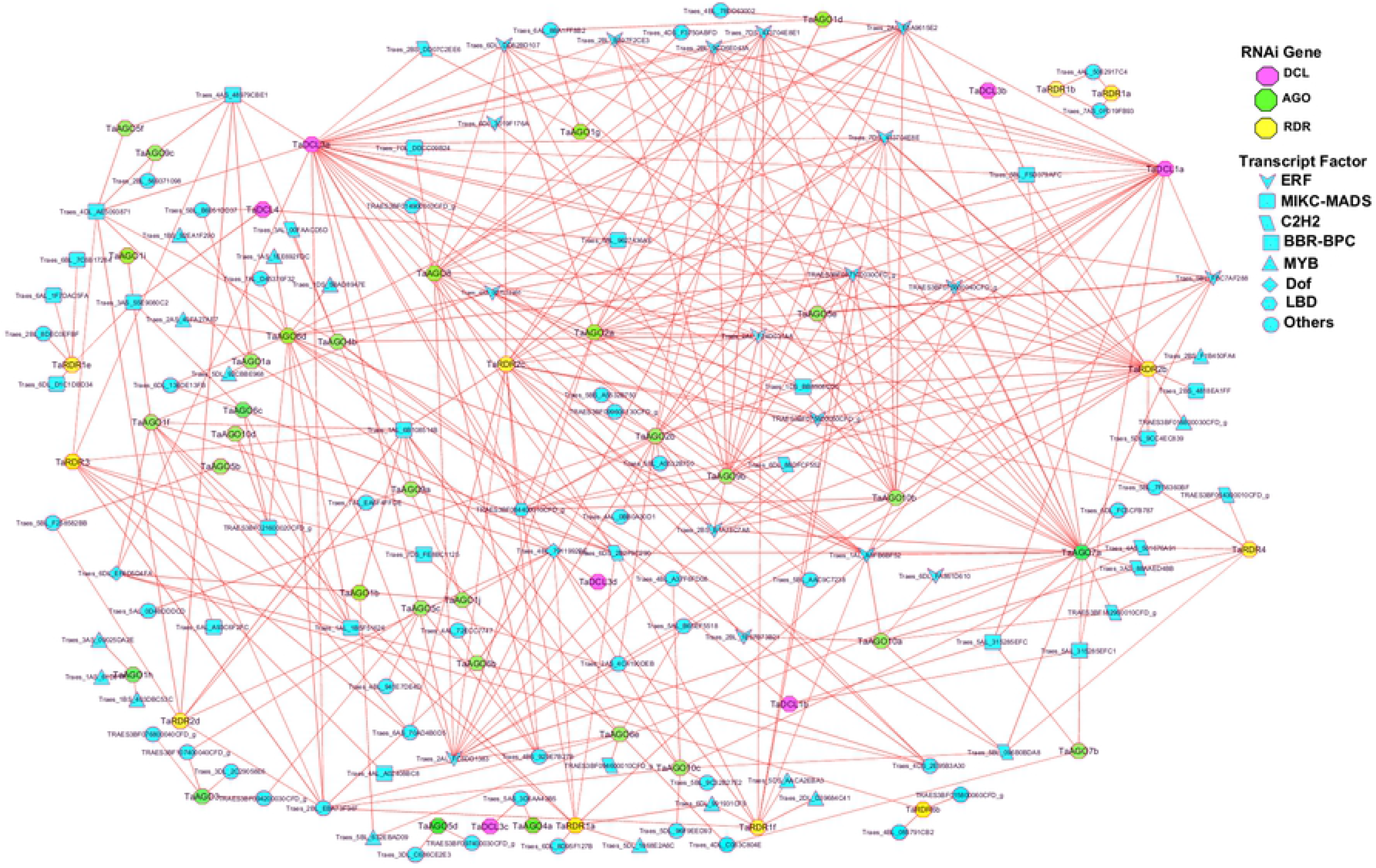
Regulatory gene network among the TF families and the predicted RNAi genes in wheat (*T.aestivum*). The nodes of the network were colored based on RNAi genes and TFs. TaDCL, TaAGO and TaRDR genes were represented by pink, green, and yellow color nodes respectively and the TFs were represented by paste color nodes. Different node symbols were used for different TF families.

Additionally, four hub TF families were selected based on the node degree criterion that possessed three or more interaction with the predicted wheat (*T.aestivum*) RNAi genes. Among them 16 TFs belong to ERF family, one TF from MIKC-MADS and C2H2, two TFs from LBD family (Fig.11E). The ERF family with accession no. Traes_2AL_E5A9615E2 regulates all 10 and a TF of ERF family with accession no Traes_5BL_F5D379AFC regulates minimum 5 hub wheat (*T. aestivum*) RNAi genes in the network followed by the two TF of family LBD with accession no.TRAES3BF084400010CFD_g and Traes_1DS_BB8508CC6 that regulate 8 and 5 RNAi genes respectively (Fig. 11E). The TF family C2H2 and MIKC-MADS each regulates 4 and 3 RNAi genes respectively in wheat (*T.aestivum*) in the network (Fig. 11E). It is however observed from the Fig. 11E that TFs from ERF family are largely bind to the hub wheat RNAi genes that suggests that TFs from this family contain high tendency to bind with RNAi genes in particular gene expression in wheat. From earlier investigation, it is also broadly well-known that transcription factors of ERF family play a significant role in regulating the growth, evolution, and response of plants to various stresses as a signal transmission pathway in plants [85]. This TF family contains DRE-binding proteins (DREBs) that stimulate the expression of abiotic stress-responsive genes through specific binding to the dehydration-responsive element/C-repeat (DRE/CRT) *cis*-acting component in their promoters[86]. Transcription factors of ethylene-responsive factor (ERF) family also possess evolving role as the plant TF for coordinating wound defense responses and repair[87].

**Fig.11.**
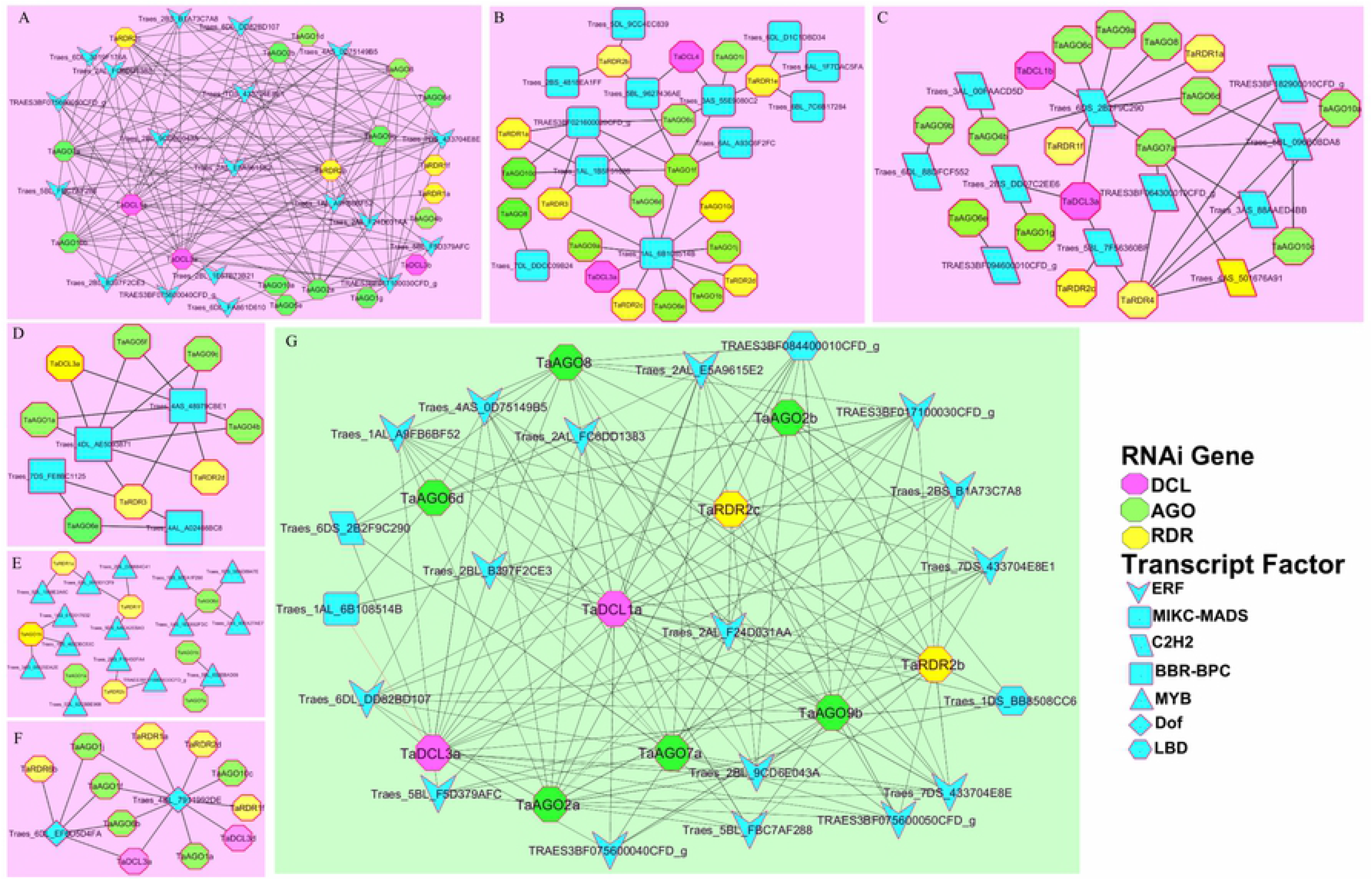
RNAi gene mediated sub-networks with different TF families. (A) ERF, (B) MIKC-MADS, (C) C2H2, (D) BBR-BPC (E) MYB (F) Dof TF family. (G) Sub-network among the hub TF families those regulate three or more RNAi genes in wheat (*T.aestivum*).

Furthermore, from this analysis (Fig. 11E), it is noticeable that, all hub TF families are associated with 10 RNA silencing machinery gens in wheat (*T. aestivum*) viz., two RNAi genes from TaDCL family (TaDCL3a and TaDCL1a), 6 RNAi genes from TaAGO family (TaAGO2a/2b, TaAGO6d, TaAGO7a, TaAGO8, and TaAGO9), and two from TaRDR family (TaRDR2b and TaRDR2c). Interestingly, only TaDCL3a member from DCL family is regulated by all four TF families (Fig. 11E) that is assumed to possess high biological importance in regulating gene expression in wheat (*T.aestivum*) plants.

### 3.8 Cis-acting regulatory element analysis in wheat (*T.aestivum*) RNAi gene families

*Cis*-regulatory elements (CREs) act as enhancers and promoters for particular gene expression in living organisms. CREs take part in indispensable activities for development and physiology through the regulation of gene expression in plants[88]. Generally these *cis*-acting components are consists of non-coding DNA possessing binding regions for TFs and/or other regulatory molecules for triggering gene transcription[88,89]. In this work, CREs analyses were also carried out to study their functional divergence in wheat (*T.aestivum*) plant for the identified RNAi genes using PlantCARE database. It is obvious from the Fig. 12 that most of the *cis*-regulatory sequences/motifs are involved in light responses (LR) which are diversely possessed by most of the RNAi gene families’ promoter sites in wheat (*T.aestivum*). Plants photosynthesis phenomenon is greatly connected to light response that mostly takes place in leaves. Photosynthesis is also the key physiological parameter in wheat plants that relates ultimately in many aspects to increase grain quality and crop productivity[90]. Increase photosynthesis rate can utilize the solar radiation properly which leads to create early flowering time because flowering signals are produced in leaves [91,92]. Therefore, the identified CREs related to LR is assumed to retain direct connections for high photosynthesis level in wheat plant leaves. Amongst the LR CREs the ATCT-motif, ATC-motif, Box-4, AE-box, G-box, I-box, GAT-motif, GT1-motif were shared by the most of the RNAi genes in wheat (*T.aestivum*) (Fig12). The TC-rich repeats (CRE involved in defense and stress responsiveness), MBS (MYB binding site involved in drought inducibility), and LTR elements (CRE involved in low-temperature responsiveness) were commonly found as stress responsive CREs among the RNAi gene families in wheat (*T.aestivum*).

**Fig 12.**
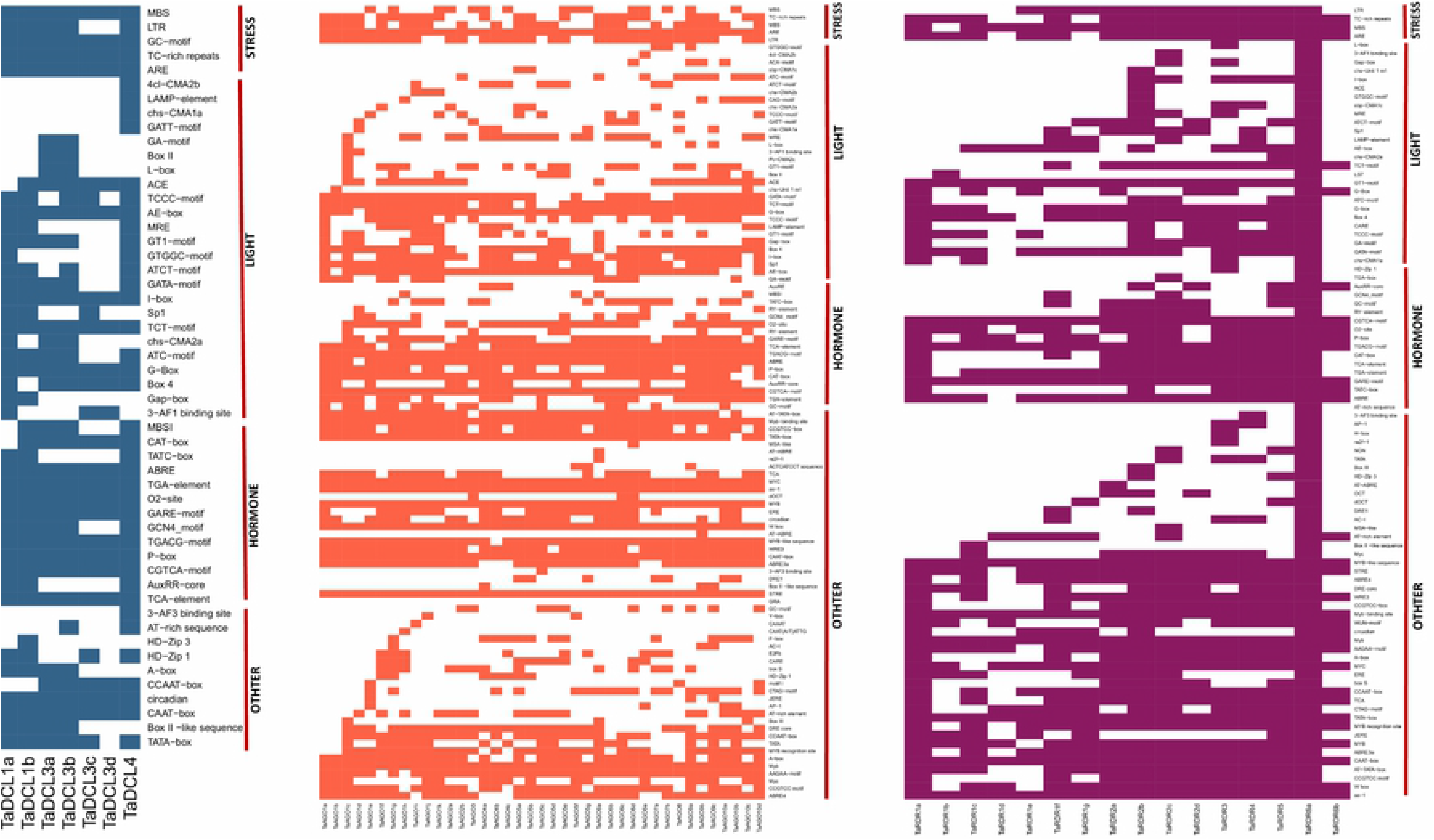
The CREs of the promoter sites in the predicted RNAi gene families of DCL, AGO and RDR in wheat (*T.aestivum*). The dark color represents the existence of a *cis*-acting element corresponding to each RNAi gene member.

It is well-studied that various plant hormones or phytohormones are essential for plant growth and developmental timing[93]. More than a dozen common CREs is related to wheat plant hormonal activities was also identified from the PlantCARE database analysis. The ABRE (*cis*-acting element involved in the abscisic acid responsiveness), AuxRR-core (*cis*-acting regulatory element involved in auxin responsiveness), GC-motif (enhancer-like element involved in anoxic specific inducibility), GARE-motif (gibberellin-responsive element), O2-site (*cis*-acting regulatory element involved in zein metabolism regulation), P-box (gibberellin-responsive element), TATC-box (*cis*-acting element involved in gibberellin-responsiveness), TCA-element (*cis*-acting element involved in salicylic acid responsiveness) and TGA-element (auxin-responsive element), RY-element (*cis*-acting regulatory element involved in seed-specific regulation), MBSI(MYB binding site involved in flavonoid biosynthetic genes regulation), GCN4_motif(*cis*-regulatory element involved in endosperm expression), CGTCA-motif and TGACG-motif (cis-acting regulatory element involved in the MeJA-responsiveness), CAT-box and NON-box(cis-acting regulatory element related to meristem specific activation)were predicted as the hormone responsive CREs in wheat (*T.aestivum*)(Fig.12). These however are retained by the three TaDCL, TaAGO and TaRDR protein families which are termed as phytohormones CREs. Previous studies also suggested that five most important plant hormones: auxin, gibberellin, cytokinin, ethylene, and abscisic acid act collectively or individually to effect plant growth and development[93–97].

Besides, there are some other CREs were also found and identified in the promoter and enhancer sites of the RNAi genes in wheat (*T.aestivum*) assumed to be responsible for a few momentous activities for controlling gene expression. The AT-rich element (binding site of AT-rich DNA binding protein (ATBP-1), AT-rich sequence (element for maximal elicitor-mediated activation),CAAT-box (common *cis*-acting element in promoter and enhancer regions), CCAAT-box (MYBHv1 binding site), TATA-box (core promoter element around-30 of transcription start), circadian (*cis-*acting regulatory element involved in circadian control) etc. were recognized as other *cis*-acting regulatory element shared by RNAi genes in wheat(*T.aestivum*) (Fig 12). In addition, some unknown CREs was discovered together with the reported *cis*-elements (S6, S7, and S8 Data). However, the identification and characterization of the predicted CREs associated with the suggested RNAi genes in wheat (*T.aestivum*) would however provide strong basis to plant molecular researchers for in-depth molecular investigation about their role in plant growth and development in diverse environmental conditions as whole.

### 3.9. *In silico* expressed sequence tag (EST) analysis for RNAi genes in wheat (*T.aestivum*)

In order to obtain further insights about the genomic information in terms of gene expression at various tissues or organs with varied conditions, computational expressed sequence tag(EST) data analysis was carried out for all 62 RNAi gene families in wheat (*T.aestivum*) using web-based PlantGDB database. This analysis results discovered that numerous RNA silencing machinery genes of DCL, AGO and RDR family showed their expression in several different important tissues and organs in wheat (*T.aestivum*). Also the expression study of the RNAi gene families of DCL, AGO and RDR groups in several plant species was investigated and thus the analyses results also suggested that RNAi gene families showed considerable and significant expression eminence in root, leaf, flower, seed, endosperm, spike and other important organs or tissues[1,6,7,10,35–38,98]. It is however observed from the Fig. 13 that all most all members of TaDCL, TaAGO and TaRDR gene families exhibited their expression at least in one tissue or organ while TaAGO5c, TaAGO6e and TaAOG7b did not show any expression in any tissue or organ in wheat(*T.aestivum*). Nearly 50% RNAi genes of all 62, expressed in root, spike, anther, heads, shoots and endosperm followed by floret, leaf and seed (Fig.13) that certainly implied that these tissues and organs provide major contribution for improved wheat grain formation resulting in increased wheat yield. Among the DCL gene members, TaDCL4 exhibited on in root, wheat head, and seedling shoot specific expression. All TaDCL3 gene sub members displayed expression in root, spike, anther, and endosperm only except TaDCL3d that was in endosperm specific (Fig.13). On the other hand, TaDCL1a and TaDCL1b had no root, leaf, wheat head/shoot and floret/flower specific expression (Fig.13) but interestingly they had only endosperm-embryo, seed, spike specific expression characteristics which indicates that these two gene members of TaDCL family might play role in formation of healthy and sufficient number of grains in wheat plant(*T.aestivum*).

**Fig.13.**
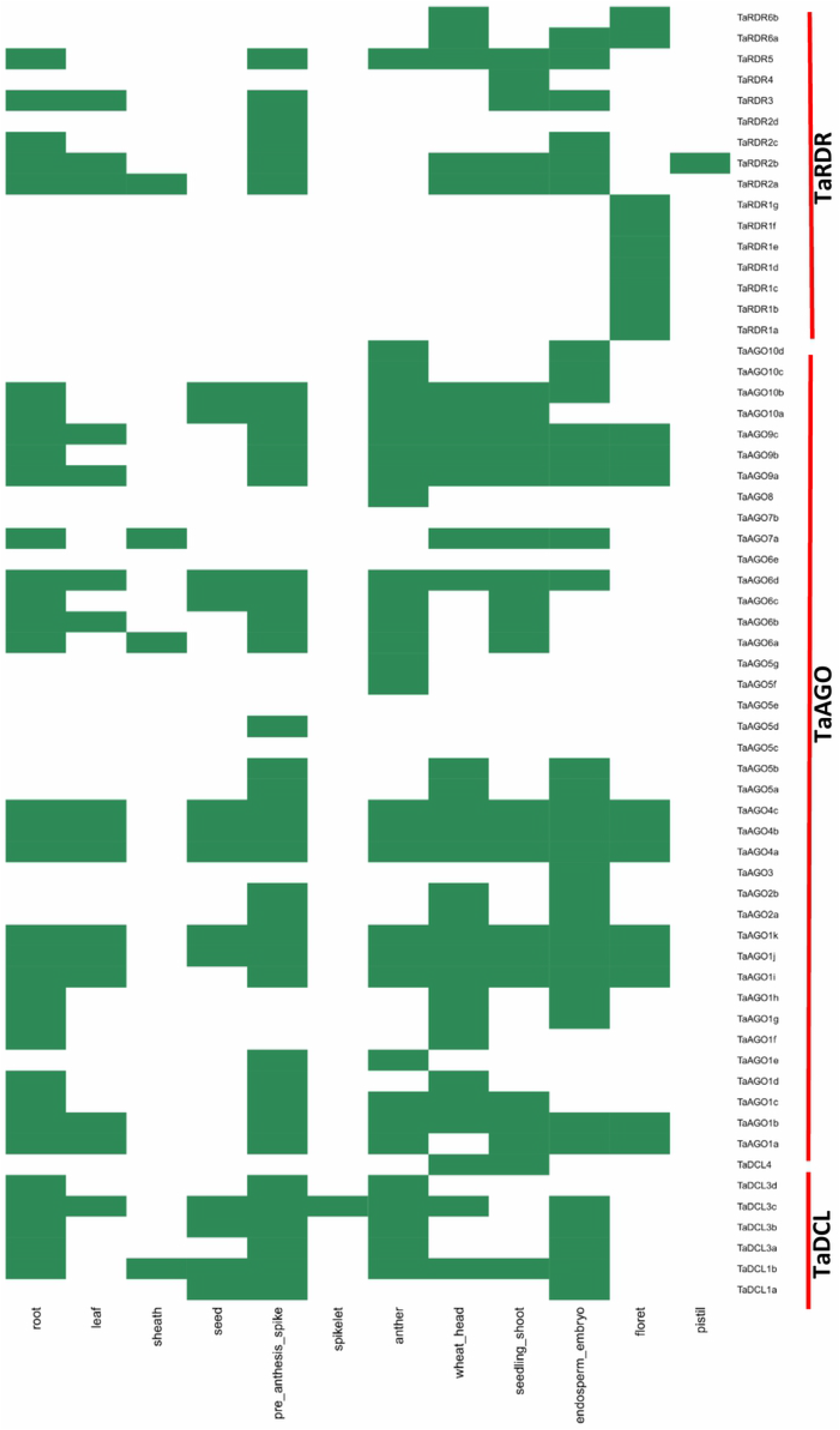
The *in silico* expressed sequence tag (EST) analysis of the identified RNAi genes in *T.aestivum*. The green color represents the existence of expression of the corresponding RNAi genes in wheat (*T. aestivum*).

Among the all TaAGO and TaRDR gene families, TaAGO1e, TaAGO3, TaAGO2a/2b, TaAGO10d/10c, all 7-sub members of TaAGO5 family, TaRDR2d, TaRDR4 and TaRDR6a/6b had no root specific expression (Fig.13). Though the TaAGO2a/2b genes expressed in wheat endosperm, head, and spike (Fig. 13) implied that these two genes contribute in grain development. But surprisingly three sub members of TaAGO1 (TaAGO1i/j/k) as well as all three sub families of TaAGO4 and TaAGO9 showed expression in all mentioned tissues and organs in the expression map except pistil, spikelet and sheath (Fig.13). This indicates that these 6 members of TaAGO4 and TaAGO9 might have roles in wheat plant growth and development. The gene sub-family TaAGO3 had only endosperm-embryo specific expression. The gene sub families TaAGO6a/6b/6c had shoot anther, spike, and root specific expression whereas the two members TaAGO6c/6d showed leaf and only TaAGO6d showed endosperm specific expression. TaAGO7a also showed root, wheat head/shoot and endosperm specific expression (Fig.13). Among the four sub families of TaAGO10 gene, the TaAGO10a/10b retained expression in all tissues and organs of wheat except leaf, sheath, and floret (Fig.13). Earlier wet lab study reported that RNAi genes of AGO family are tend to show diverse expression intensity in leaf, stem, flower in *Brassica napus*, pepper(*Capsicum Annuum*)[1,39].

Furthermore, EST analysis for all sub members of TaRDR1 showed their expression in flower but had no expression in any tissues and organs in wheat (*T.aestivum*) (Fig.12). It that suggests that these sub gene families are tend to responsible for expression in flower like two sub families TaRDR6a/6b(Fig. 13).Previous study on pepper(*Capsicum Annuum*) suggested that large number of RDR gene families showed flower specific expression[1]. In contrast, four sub members of TaRDR2 family did not provide any expression in flower. However, all members of TaRDR2 showed expression all most in all mentioned important tissues and organs with additional expression of TaRDR2b in pistil in wheat (*T.aestivum*) but TaRDR2d expressed in only spike (Fig. 12). It can be said from this EST analysis of all 62 RNAi genes in wheat (*T.aestivum*) that genes those expressed in root and leaf might play role in resistance to abiotic stresses such as drought and water logging etc. Genes expressed in leaves may contribute in promoting photosynthesis rate to produce sufficient energy for the wheat plants. Increasing photosynthesis level stimulate the flowering time of the crop as the respective signals are produced in leaves[91,92].Seed and endosperm specific expression of the RNAi genes demonstrates that they might have significant role in maintaining quality seed formation.

### 3.10. Expression analysis of TaDCL Genes in leaves, roots and against drought Stress

To obtain evidence for the likely functions of the TaDCL genes, their expression profile in leaves and roots of two-week aged plants were analyzed by qRT-PCR. The results showed that two out of seven TaDCL genes (TaDCL3a and TaDCL3b) expressed significantly in root and had no expression in leaves indicating root specific function of TaDC3a and TaDCL3b gene families that validates the respective EST analysis too (Fig. 14A and Fig.13). To explore the role of TaDCL genes in response to drought stress tolerance, the expression patterns of TaDCLs were measured at 3-days post treatment against drought. The results showed that drought treatment positively induced the expression of all the seven TaDCL candidate genes. Among them, TaDCL3b and TaDCL4 showed the significant change in expression in response to drought signifying these genes might play a vital role against drought stress in wheat(*T.aestivum*) (Fig. 14B). The results from this analysis however can enrich our understanding of DCL genes in wheat and constitute a base for advance study.

**Fig.14.**
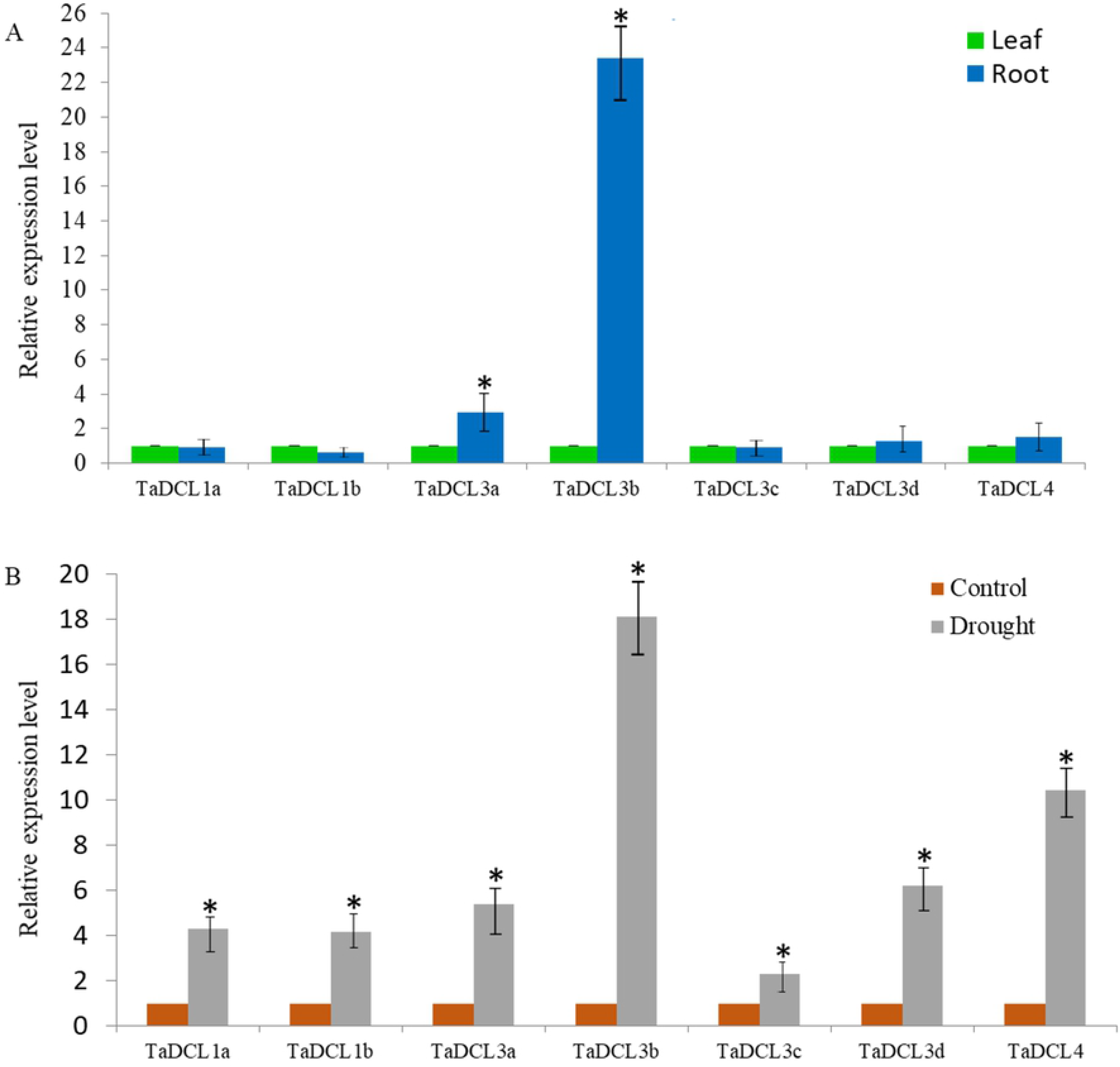
qRT-PCR expression of TaDCL genes in (A) leaves and roots and (B) under drought stress. *T.aestivum* 18s was considered as the reference gene, and three biological replicates were accomplished for the experiments. Error bars indicate the standard error. Asterisks indicate the significance difference between control and treatment at p-value < 0.05

## 4. Conclusion

Wheat as a cereal crop is consumed by a large community of the people across the globe for the key source of carbohydrates after rice. Due to of its high protein content, this crop is also used for producing many important human foods for each day. Genome-wide identification, characterization, and diverse functional analyses of the RNA silencing machinery gene families were top most priority because these set of gene families carry great significance for regulating gene expression by adopting gene-silencing mechanism in plants by restricting the transcript buildup in cells. In this work, a number of bioinformatic techniques were applied for comprehensive identification, characterization, as well as *in-silico* biological and molecular functional activities were predicted in wheat (*T.aestivum*). Analyses results suggest that the wheat genome possessed 7 DCL, 39 AGO and 16 RDR RNAi genes. Initial analysis provided some standard genomic and physicochemical info for the identified genes and corresponding proteins. The phylogenetic analysis provided that all subfamilies of these three gene sets maintain their evolutionary relationships similar to their rice and *Arabidopsis* homologs. First subfamily of DCL, AGO and RDR possessed the multiple copies of genes higher than that of corresponding rice and *Arabidopsis.* Conserved functional domain and motif structure analyses suggested that these genes also contained consistent domain and motif structure similar to those of rice and *Arabidopsis* RNAi genes. Besides, GESA and SCL analyses established the insights into the functional routes and their involvement scenario in RNAi process in wheat (*T.aestivum*).We discovered the most important regulatory relationship network amongst the popular TF families and the putative RNAi gene families. Promoter and enhancer as CREs was analyzed those also attached with TFs to control the gene expression for some specific reasons during the wheat plant’s growth and development. CRE and EST analyses hence suggests that the identified TaDCL, TaAGO and TaRDR gene families have diversifying participation for plant growth, development, maintenance the grain quality and increased crop production in wheat(*T.aestivum*). Wet lab expression study demonstrated that only TaDCL3a and TaDCL3b had root specific expression that validates EST analysis as well. Finally, all TaDCL transcripts induced up-regulation in drought stress condition with high expression in TaDCL3b and TaDCL4. The results of this study thus provide the important indication of evolutionary resemblances of DCL, AGO and RDR genes with their rice and *Arabidopsis* counterpart. Obtained results would however provide valuable sources for further biological and molecular justification and implementation to draw more specific conclusion regarding any particular gene(s) and its domain of activity against different biotic and abiotic stresses as well as growth and improvement in wheat plants for more improvement of this potential cereal crop across the world.

## Acknowledgments

We are very grateful to Bangladesh Agricultural Research Institute and Bioinformatics Lab, Department of Statistics, University of Rajshahi to provide us opportunity to conduct this research using lab facilities. MA of icddr,b thanks the Governments of Bangladesh, Canada, Sweden, and the United Kingdom for core/unrestricted support.

## Abbreviations

RNA: Ribonucleic Acid
RNAi: RNA interference
siRNA: Short interfering RNA
miRNA: micro RNA
ssRNA: Single-stranded RNA
dsRNA: Double-stranded RNA
DCL: Dicer-like
AGO: Argonaute
RDR: RNA-dependent RNA polymerase
RISC: RNA-induced silencing complexes
DRM: Double-stranded RNA-binding motif
BLAST: Basic Local Alignment Search Tool
MEGA: Molecular Evolutionary Genetics Analysis
GO: Gene Ontology
SCL: Sub Cellular Location
NJ: neighbor-joining
PAZ: Piwi Argonaut and Zwille
DUF: Domain of unknown function
GO: Gene Ontology
GSDS: Gene Structure Display Server
HMM: Hidden Markov Model
MEME: Multiple EM for Motif Elicitation
PTGS: post-transcriptional gene silencing
TF: transcription factor
CRE: *cis*-regulatory elements
qRT-PCR: quantitative reverse transcription polymerase chain reaction

## Author’s contributions

ZA and MM designed the research. ZA analyzed the data using bioinformatics and statistical tools as well as drafted the manuscript. ZA and HR carried out the gene expression study. MAA performed GO and SCL analysis. MA, MJA, and MPM attended at the meeting regarding this manuscript. All authors read and approved the final version of the manuscript.

## Supporting information

**S1 Table. Primer sequences of seven TaDCL genes for qRT-PCR analysis**

**S2 Table. Percentage of three groups of RNA silencing genes involved in different cellular location in wheat (*T.aestivum*)**

**S3 Table. Distribution of TF families those regulating RNAi genes in wheat (*T.aestivum*)**

**S1 Data.** Protein sequences of the identified DCL genes in wheat(*Triticum aestivum*).

**S2 Data.** Protein sequences of the identified AGO genes in wheat(*Triticum aestivum*).

**S3 Data.** Protein sequences of the identified RDR genes in wheat(*Triticum aestivum*).

**S4 Data.** GO enrichment analysis result for reported RNAi genes.

**S5 Data.** List of transcript factors and their families regulating the predicted RNAi genes.

**S6 Data.** List of *cis*-regulatory element associated with the TaDCL protein families

**S7 Data.** List of *cis*-regulatory element associated with the TaAGO protein families

**S8 Data.** List of *cis-*regulatory element associated with the TaRDR protein families

**S1 Fig.** GO slim hierarchy tree analysis of the predicted RNAi genes in wheat (*T.aestivum*) was projected using QuickGO web-based bioinformatic tool.(A) Biological process(BP) (B) Molecular Function(MF) and (C) Cellular Component(CC). GO IDs are highlighted in upper part of the boxes shaded blue color and their corresponding funcitonal terms are mentioned in the middle of each box. Boxes with ligh yellow color are statistically significant GO functions whereas the boxes with white color are insignificant GO.

**S2 Fig.** Distribution of TF families corresponding to each RNAi gene member. Rows of the figure represent the predicted RNAi genes and the columns represent the TF families. The number inside the cell indicates the member of TFs of the corresponding family involve in regulating the RNAi genes in wheat(*T.aestivum*).

